# Chronic administration of XBD173 ameliorates cognitive deficits and neuropathology via 18 kDa translocator protein (TSPO) in a mouse model of Alzheimer’s disease

**DOI:** 10.1101/2023.02.23.529740

**Authors:** Arpit Kumar Pradhan, Tatjana Neumüller, Claudia Klug, Severin Fuchs, Martin Schlegel, Markus Ballmann, Katharina Johanna Tartler, Antoine Pianos, Maria Garcia-Sanchez, Philippe Liere, Michael Schumacher, Matthias Kreutzer, Rainer Rupprecht, Gerhard Rammes

## Abstract

Alzheimer’s disease (AD) is characterized by the accumulation of β-amyloid peptide (Aβ). There is increasing evidence that depression may precede AD and may be an early manifestation of dementia, suggesting common mechanisms underlying both diseases. Ligands targeting the mitochondrial translocator protein (18 kDa) (TSPO), promote neurosteroidogenesis and may be neuroprotective. Moreover, TSPO is upregulated in AD. To study whether the TSPO ligand XBD173 may exert early neuroprotective effects in AD pathology we investigated the impact of XBD173 on amyloid toxicity and neuroplasticity in mouse models. We show that XBD173 (emapunil), via neurosteroid-mediated signaling via delta subunit-containing GABA_A_ receptors, prevents the neurotoxic effect of Aβ on long-term potentiation (CA1-LTP) in the hippocampus and prevents the loss of spines. Chronic but not acute administration of XBD173 ameliorates spatial learning deficits in transgenic AD mice with arctic mutation (ArcAβ) mice. The heterozygous TSPO-knockout crossed with the transgenic arctic mutation model of AD mice (het TSPOKO X ArcAβ) treated with XBD173 does not show this improvement in spatial learning suggesting TSPO is needed for procognitive effects of XBD173. The neuroprotective profile of XBD173 in AD pathology is further supported by a reduction in plaques and soluble Aβ levels in the cortex, increased synthesis of neurosteroids, rescued spine density, reduction of complement protein C1q deposits, and reduced astrocytic phagocytosis of functional synapses both in the hippocampus and cortex. Our findings suggest that XBD173 may exert therapeutic effects via TSPO in a mouse model of AD.

## Introduction

Alzheimer’s disease (AD) is characterized by memory impairment and a progressive cognitive decline which has been attributed to the accumulation of Aβ in the brain. AD pathophysiology is largely reflected by the extracellular aggregation of Aβ peptide, hyperphosphorylated tau aggregates, and reactive gliosis (1). Increased levels of pro-inflammatory cytokines such as interleukin-1b (IL-1b) have been shown to promote neurotoxicity induced by N-methyl-D-aspartate (NMDA) receptors, which is further amplified by the presence of Aβ (2, 3). Aβ, a product of proteolytic cleavage (by β-secretases and γ-secretases) of amyloid precursor protein (APP) via the amyloidogenic pathway, may accumulate in AD brains both as a consequence of excessive production and impaired clearance (4). The Aβ_1–42_ peptide has a higher propensity to aggregate and is highly enriched in amyloid deposits. Meanwhile, numerous studies have reported that soluble Aβ_1-42_ oligomers impair cognitive function and inhibit long-term potentiation (LTP), a cellular correlate for learning and memory (5–8). Ample evidence suggests that Aβ_1-42_ pathologic effects are mediated by alterations in glutamatergic signaling, specifically NMDA receptors and possibly metabotropic glutamate receptors (9).

Chronic depression has been considered an antecedent of AD and may constitute an early manifestation of dementia before the cognitive decline becomes apparent (10–12). This suggests that depression may be a risk factor for AD and calls for research that addresses the relationship between both diseases. As such, novel compounds that target both depression and dementia would constitute a novel therapeutic approach. The 18 kDa TSPO protein, located on the outer mitochondrial membrane, is known to orchestrate several important functions including cholesterol transfer into mitochondria, TSPO ligands-induced neurosteroidogenesis, mitochondrial homeostasis, gliosis, and apoptosis (13, 14). Contrary to benzodiazepines which are positive allosteric modulators of GABA_A_ receptors, neurosteroids act through a different site than that targeted by benzodiazepines. For example, the δ-subunit of GABA_A_ receptor is considered to be highly sensitive to neurosteroids but not to benzodiazepines (15). This potentially explains the positive effects of synaptic modulation by neurosteroids in view of a more favorable side-effect profile. Interestingly, in AD, TSPO upregulation in astrocytes has been observed prior to the upregulation in microglia and is considered to be an exclusive marker of glial activity in AD (16).

In this study, we therefore asked whether XBD173, a TSPO ligand known to exert an anxiolytic effect both in rodents and in humans (13, 17), may confer neuroprotective benefits in a murine model of AD. We show that XBD173 promotes TSPO mediated synthesis of different neurosteroids which, upon release, elevates GABA_A_ receptor activity containing GABA delta subunit and prevents spine loss caused by the Aβ_1-42_ oligomer. Importantly, chronic intraperitoneal injection of XBD173 reversed the spatial learning deficits in the rodent AD model. However, neuroprotection by XBD173 was absent in the het TSPOKO X ArcAβ animals, thus confirming the role of TSPO in XBD173 mediated neuroprotection. This is further supported by a reduction in plaques, soluble Aβ levels in the cortex, increased synthesis of neurosteroids, rescued spine density, and reduced astrocytic phagocytosis of functional synapses both in the hippocampus and cortex. Moreover, we show that the activation of TSPO impairs the complement pathway protein C1q and reduces the hyperactivity of the complement system.

## Materials and Methods

### Animals

All animal study protocols were in accordance with the German law on animal experimentation and were approved by the animal care committee (Technical University Munich, Munich, Germany) and the experiments performed confirmed the institutional regulatory standards. Mice (maximum of 6 mice per cage) were caged in a climate-controlled room (23 ± 0.5 °C) with 12 hours of light and darkness with an *ad libitum* supply of food and water. All the acute slice experiments involved mice of both sexes which were 8-12 weeks old. Animals involved in the behavioral testing were housed under similar conditions except for two weeks before the water cross maze behavioral testing where mice were housed in a reverse light-dark condition (12-hour light and darkness) cabinet maintained at a temperature of 23±0.5°C. Both male and female animals at an age of 8-9 months were given either chronic or acute treatment before the behavioral experiment.

Wild-type (C57BL/6, WT) mice were obtained from Charles River (Italy) and ArcAβ (APP E693G) mice were obtained from CALCO (Italy). The transgenic mice carrying this arctic mutation (ArcAβ) present a point mutation where glutamic acid (E) has been replaced by site-directed mutagenesis with glycine (G). The TSPOKO mice were bred by our group in Munich and had a global knockout of TSPO. TSPOKO X ArcAβ mice were generated by crossing stable lines of TSPOKO and TSPOKO X ArcAβ mice. The GABA δ knock-out (δ-KO) (line B6.129-Gabrdtm1Geh/J) mouse line was bred by our group in Munich (Germany) in a way similar to the Jackson Laboratory (Jax stock 003725). As a result of the KO mutation, the δ subunit of the GABA_A_ receptor is absent in these animals. The absence of the δ subunit in the GABA_A_ receptor of these mice leads to attenuated sensitivity to neurosteroids as described previously by Mihalek et al. (1999) (18). Genotyping was done in each case to confirm the respective transgenic lines.

### Preparation of oligomeric Aβ and test compounds

Aβ_1-42_ (order number H-1368; Bachem, Bubendorf, Switzerland) and Aβ_1-40_ (Product Number: 4014442; Bachem, Bubendorf, Switzerland) were suspended in 100% hexafluoroisopropanol (HFIP, Sigma Aldrich) to a concentration of about 1 mg/400 µL HFIP and shaken at 37 ℃ for 1.5 h. Aliquots of 100 µg Aβ_1-42_ and Aβ_1-40_ were made per tube. The HFIP was then evaporated by using a Speedvac for 30 min. After the tubes were completely dry, they were labeled and stored at -80 ℃. The respective Aβ were dissolved in DMSO (Sigma Aldrich) to obtain a concentration of 100 µM. This solution was then diluted to a concentration of 50 nM Aβ_1-42_ and Aβ_1-40_ using Ringer’s solution.

Diazepam and XBD173 were dissolved in DMSO. For LTP recordings, XBD173 was dissolved in aCSF to obtain a 300 nM concentration whereas, for other *ex vivo* slice experiments, XBD 173 was dissolved to a 3 µM concentration. For chronic and acute treatment for behavioral experiments, XBD173 and diazepam were diluted in a 0.9% NaCl solution. Drug doses were 1mg/kg for XBD173 and 1mg/kg for diazepam. For chronic treatment, intraperitoneal (i.p.) injections of XBD173, diazepam, or vehicle were given every alternate day for 12 weeks starting from 8-9 months while for the acute treatment group, the treatment XBD173(1 mg/kg), and diazepam (1mg/kg) was given once 3 days before starting of the behavioral tests. The vehicle group received only DMSO diluted in 0.9 % NaCl solution at a similar concentration as with XBD173.

### Acute slice preparation

Mice were deeply anesthetized with isoflurane before decapitation. The mouse brain was rapidly placed in an ice-cold Ringer solution consisting (in mM) of 125 NaCl, 2.5 KCl, 25 NaHCO_3_, 0.5 CaCl_2_, 6 MgCl_2_, 25 D-glucose and 1.2 NaH_2_PO_4_, with a pH of 7.3 and continuously oxygenated with carbogen gas (95% O_2_/5% CO_2_). Sagittal hippocampal slices (350 µm thickness) were obtained from both hemispheres using a microtome (HM 650 V; Microm International, Germany). Slices were recovered for 30 min at 35 °C in a chamber submerged with artificial cerebrospinal fluid (aCSF) containing (in mM) 125 NaCl, 2.5 KCl, 25 NaHCO_3_, 2 CaCl_2_, 1 MgCl_2_, 25 D-glucose and 1.2 NaH_2_PO_4_, and constantly oxygenated with carbogen. The slices were then placed at room temperature for 1 hour of recovery before transferring them into the recording chamber. The slices in the recording chamber were fixed using a platinum ring with two nylon threads while carbonated aCSF was steadily perfused at a flow rate of 5 ml min^-1^. All recordings were performed at room temperature (21-23 °C).

### CA1-LTP recordings

LTP recordings were performed as previously established (19). In brief, we recorded field excitatory postsynaptic potentials (fEPSPs) using glass micropipettes (1–2 MΩ) filled with aCSF solution in the CA1 region of the hippocampus. fEPSPs were recorded in the hippocampal CA1 stratum radiatum induced by stimulation in the Schaffer collaterals-commissural pathway. fEPSPs were evoked via alternating test stimulation (50µs, 5-20V) delivered through bipolar electrodes (Hugo Sachs Elektronik-Harvard Apparatus, Germany; 50µm tip diameter) placed on both sides of the recording pipette. In this way, the non-overlapping populations of fibers of the Schaffer collateral-associational commissural pathway were stimulated. The stimulation intensity was adapted to values evoking an fEPSP slope of around 25-30% of the maximum response, for baseline recording. LTP was induced after at least 20 minutes of recording a stable baseline by delivering a high-frequency stimulation (HFS) train (100 pulses at 100Hz over 1s) through one of the two stimulating electrodes. The independent stimulation with two electrodes allowed the recording of internal control in the same slice. Following the administration of HFS in the absence of any substance, the potentiation of the fEPSP slope was measured for 60 minutes after the tetanic stimulus, while maintaining the settings used for the baseline. Briefly, HFS was delivered from one of the electrodes in the absence of Aβ (Aβ_1-40_ or Aβ_1-42_), and potentiation of the responses was monitored for at least 60 minutes following tetanus. Aβ (Aβ_1-40_ or Aβ_1-42_) was then applied via the bath solution for 90 minutes, before inducing the LTP in the second input via the HFS train.

To test toxicity prevention, HFS was delivered from one of the electrodes in the presence of either XBD173, pregnenolone, allopregnanolone, or allotetrahydrodeoxycorticosterone (THDOC), and the potentiation of the responses was monitored for at least 60 minutes after tetanic stimulation. After that, Aβ (Aβ_1-40_ or Aβ_1- 42_) was applied via bath solution for 90 minutes before inducing LTP in the second input via the HFS train. LTP inhibition or blockage was defined as the fEPSP slope being less than 120% of the prestimulation slope in the last 10 minutes after HFS. The fEPSPs were amplified (BA-2S, npi electronic, Germany), filtered (3 kHz), and digitized (9 kHz) using an interface board (ITC-16, Instrutech Corp., USA) and the WinLTP program. Because stimuli were alternately delivered to each input every 15 seconds, two signals from the same input were averaged to one for analysis and one data point per minute. WinLTP program was used to analyze the fEPSPs offline. The slope of the fEPSP was measured between 20% and 80% of the peak amplitude which was then normalized to the 20-minute control baseline recording obtained before the induction of LTP.

### Spine imaging

We used Thy1-eGFP mice for the dendritic spine analysis of *ex vivo* experiments with Aβ_1-42_. After the recovery period of 1 hour at room temperature, the slices were treated with XBD173 and Aβ_1-42_. After incubation with respective treatment, slices were collected, fixed with 4% PFA for 2 days, and cryoprotected with 30% sucrose for 3 days. A cryostat (CryoStar NX70, Thermo Fisher Scientific, Bremen, and Germany) was used to prepare slices at a 100 µm thickness (Bremen, Germany). The slices were transferred to slides and coverslipped after being washed with PBS. Images of apical oblique dendritic spines in the CA1 region of the hippocampus were captured with a confocal microscope (Leica SP8, Wetzlar, Germany) with a Z step size of 0.3 µm and a 60X/1.40 N.A. oil-immersion objective. 6-8 dendrites of 20 - 40 µm in length from pyramidal neurons were obtained from the stratum radiatum layer in the CA1 region per mouse and were analyzed with the IMARIS 9.7 for Neuroscientists (Oxford Instruments, bitplane). Reconstruction of dendrites and spines was also performed using the Filament tool from the IMARIS package before categorizing the spines into individual subtypes. Dendritic spines were categorized as thin, mushroom, and stubby subtypes, based on established criteria. The following parameters were applied to automatically categorize the spines in the IMARIS: Mushroom Spines: length (spine) < 3 and max_width (head) > mean_width (neck) * 2; Long Thin Spines: mean width(head) >= mean_width(neck) and Stubby Spines: length(spine) < 1. The spine density was defined as the number of spines per µM of the dendrite.

### Flow cytometry

After pharmacological treatment with XBD173 and Aβ_1-42_, the hippocampus and cortex were separated for each slice. These were then transferred to a 1 ml digestion solution (activated at 37 °C for 5 min) consisting of ethylenediaminetetraacetic acid (EDTA) (2 mM), L-cysteine (5 mM), and papain (5 units/ml). The enzymatic digestion was performed for 15 min at 37 °C. After the dissociation of slices, the suspension was filtered through a 70 µm cell strainer to obtain a single-cell suspension. After rinsing the cell strainer with 2 ml PBS, the samples were centrifuged for 10 minutes at 600 x g at room temperature. The supernatant was discarded and the cell pellet was resuspended in 1 ml Fluorescence-activated cell sorting (FACS) buffer consisting of EDTA (1 mM), sodium azide (15 mM), and 1% w/v bovine serum albumin (BSA). Cell count and % viability per sample were checked via a hemocytometer. To prepare the cells for FACS analysis, they were spun down at 600 x g for 10 minutes at 4 °C before staining with fluorescent antibodies. The cell pellets were resuspended in a 50 µl blocking mix (49 µl FACS buffer + 1 µl anti-CD16/32 antibody) per sample and incubated in the dark for 5 minutes at 4 °C. The samples were then incubated in the dark at 4 °C for 30 minutes with 11.5 µl of the antibody full-staining mix (**Table 1**). After adding 100 µl of FACS buffer, the cells were centrifuged at 600 x g for 5 minutes at 4 °C. Afterwards, the pellets were resuspended in 150 µl of FACS fixation buffer and stored at 4 °C until the FACS analysis was performed. FACS analysis was performed by the BD LSRFortessa™ X-20 using the FACSDiva™ data acquisition software. Compensation was performed for each channel using compensation data measured previously with compensation beads. The events were then gated for single microglial cells. **Supplementary Figure 4** depicts the gating strategy. Antibodies used in the FACS analysis and the source are listed in **Table 2**.

**Table 1.**

| Antibody target | Coupled fluorochrome | µl per sample |
| --- | --- | --- |
| CD45 | FITC | 1 |
| CD11b | APC-Cy7 | 1 |
| F4/80 | PerCP-Cy5.5 | 1.5 |
| TMEM119 | PE-Cy7 | 2 |
| P2RY12 | APC | 2 |
| CD163 | PE | 2 |
| CD80 | BV605 | 1 |
| CX3CR1 | BV421 | 1 |
|  |  | 11.5 |

**Table 2.**

| Antibodies | Catalog number | Manufacturer |
| --- | --- | --- |
| TruStain FcX™ (anti-mouse CD16/32) Antibody | 101320 | BioLegend, San Diego (USA) |
| FITC anti-mouse CD45 Antibody | 147709 | BioLegend, San Diego (USA) |
| APC/Cyanine7 anti-mouse/human CD11b Antibody | 101225 | BioLegend, San Diego (USA) |
| PE anti-mouse CD163 Antibody | 155307 | BioLegend, San Diego (USA) |
| Brilliant Violet 605™ anti-mouse CD80 Antibody | 104729 | BioLegend, San Diego (USA) |
| PerCP/Cyanine5.5 anti-mouse F4/80 Antibody | 123127 | BioLegend, San Diego (USA) |
| APC anti-P2RY12 Antibody | 848005 | BioLegend, San Diego (USA) |
| Brilliant Violet 421™ anti-mouse CX3CR1 Antibody | 149023 | BioLegend, San Diego (USA) |
| Tmem119 Monoclonal Antibody (V3RT1GOsz), PE-Cyanine7 | 25-6119-80 | ThermoFisher Invitrogen |
| Iba1/AIF-1 (E4O4W) XP® Rabbit mAb | 17198 | Cell Signaling Technology, Frankfurt am Main (GER) |

### Behavioral test

Mice were trained in the water cross maze (WCM) using a previously described hippocampus-dependent place learning protocol (20). The WCM set up tests for spatial learning and memory, where mice learn to use spatial cues to find a hidden platform to escape the water. Briefly, a plus-shaped maze made out of clear transparent plexiglass with four identical arms (length and width 50 cm and 10 cm) was designated with cardinal directions (N-, S-, E-, and W- arms). The maze was filled with fresh tap water (22 °C ± 1 °C) up to a height of 11 cm and a platform of the height of 10 cm and surface area (8 cm X 8 cm) was placed in the W arm. The light conditions were kept at < 15 lux. A divider was placed to block off the fourth arm so that the mice chooses either the E or W arm for escape. The entire period of this behavioral experiment was divided into 3 phases, a. Training phase (for a consecutive 5 days period) b. Retest 1 (performed after a week from the last day of the training phase) c. Retest 2 (performed after a month from retest 1) (illustrated in the schematics in **Figure 2 A**). Retest 1 and retest 2 were designed to measure the retention of memory from the training phase and to observe if the drug effect is short-lived or long-lived. Each day had 6 randomized trials per mouse where the starting arms were alternated in a pseudorandomized order to make sure the mouse does not get accustomed to body turn to find the platform. After every trial, the mice were kept under infrared light and allowed sufficient time for rest before the next trial. A maximum time of 30 s was allotted for each mouse to find the platform. In the event of the mouse not finding the platform, the experimenter gently guided the mouse to the platform (in such cases latency of 31 s was noted).

We used the following parameters to evaluate spatial learning and memory: 1. Escape Latency: time to reach the platform from the starting arm (averaged over 6 trials every day per mouse) 2. Accuracy: Represents the number of correct trials per day in which the mice directly swam to the correct arm containing the platform without returning to the starting arm or the false arm (with no platform) 3. Accurate Learners: Number of mice that performed at least 5 out of 6 trials accurately per day, expressed as a percentage of the total number of mice per group. Accuracy change or escape latency change from the training phase in retests was calculated by subtracting the performance in respective retests with the mean of day 4 and day 5 in the training phase.

### Methoxy staining

Each mouse was anesthetized in an isoflurane chamber with a 5 % isoflurane concentration at a flow rate of 2 L/min and the brain was decapitated using a guillotine. The brain was quickly frozen in dry ice and stored at -80 °C until further use for biochemical analyses. One hemisphere of the brain was cut in the sagittal plane into 30 µm thick slices using a cryotome (CryoStar NX70, Thermofisher). These 30 µm slices were incubated with methoxy-X04 solution (0.004 mg/ml, Tocris Bioscience, 1:1 ethanol-PBS solution) at room temperature for 30 minutes. The unspecific dye was removed by washing it three times with a 1:1 Ethanol-PBS solution and two times with Milli-Q water, and the slices were then mounted on microscope slides with a fluorescent mounting medium (Dako, Germany). 3D image stacks were obtained on an epi-fluorescence microscope (Axio Imager.M2 with ApoTome.2, Jena, Zeiss, Germany). Tile scan mode was used to image the brain slices which also allows the automatic stitching of the different brain segments. The excitation wavelength for the methoxy staining was fixed at 405 nm, and the emitted light was collected from 410 to 585 nm. The area and number of plaques were counted after the removal of the background using Fiji (ImageJ).

### ELISA

The other hemisphere of the frozen mouse brain was cut coronally and segmented to the cortex and hippocampus. The mass of the hippocampus and the cortex in the Eppendorf were noted and 8 X mass cold 5M guanidine HCl/50 mM Tris HCl was added to each sample. The samples were ground thoroughly using a pestle. The homogenate was mixed with a 3D shaker for 3 h at room temperature. The homogenates were then diluted in 10 volumes of ice-cold casein buffer consisting of: 0.25 % casein, 0.05% sodium azide, 20 µg/ml aprotinin, 5 mmol/l EDTA, 10 µg/ml leupeptin in PBS (pH 8.0). This was then centrifuged at 12,000 X g for 20 min at 4°C. The supernatant was decanted and stored on ice until it was used with the ELISA kit (Human Aß Amyloid (42) ELISA kit; Invitrogen KHB 3441). The ELISA protocol was performed per the manufacturer’s instructions. The absorbance was read at 450 nm (Tecan Sunrise; Tecan Trading AG, Switzerland) within 30 min after adding the stop solution, and the concentration was read from the standard curve.

### Immunofluorescent staining Synaptic engulfment measurement

Sagittal brain slices, 30 µm thick were used for immunofluorescent staining. The slide containing the tissue sections was fixated in 1:1 acetone isopropanol solution at room temperature for 20 minutes. The sections were then permeabilized with 0.3 % triton-X in PBS containing 0.1% tween 20 (PBS/T). This was followed by a 3X wash with PBS/T. The sections were blocked with 10% v/v normal goat serum (NGS) for 1 hour at room temperature. This was followed subsequently by a 3X wash with PBS/T. The sections were incubated overnight at 4 °C with primary antibody diluted to respective concentrations in PBS/T. The primary antibody used were GFAP (GA5) Mouse mAb (1:600; Cell Signalling Tech, 3670S), Rabbit anti- C1q antibody (1:250; Abcam, ab182451) and Rabbit anti-Synaptophysin antibody (1:600; Abcam, ab52636). On the subsequent day, the sections were incubated with secondary antibodies Goat Anti-Mouse IgG H&L (Alexa Fluor® 647) (1:700; Abcam, ab150115) and Goat Anti-Rabbit IgG Fc (Alexa Fluor® 488) (1:200; Abcam, ab150089) for 2 hours at room temperature. For nuclear staining, ProLong™ Glass Antifade Mountant with NucBlue™ Stain (Sigma Aldrich, product no. 57-50-1) was used. If nuclear staining was not necessary the slides were mounted with DAKO fluorescent mounting media (DAKO, s3023).

### Plaque Astrocyte Interaction

The slides containing sagittal brain sections (30 µm) were fixed with 4% paraformaldehyde (PFA) for 15 minutes at room temperature. The permeabilization and incubation with the primary and secondary antibodies were similar as described earlier in synaptic engulfment measurement. The primary antibody used were GFAP (GA5) Mouse mAb (1:600; Cell Signalling Tech, 3670S) and rabbit anti-C1q antibody (1:250; Abcam, ab182451). The secondary antibody used were Goat Anti-Mouse IgG H&L (Alexa Fluor® 647) (1:700; Abcam, ab150115) and Donkey anti-Rabbit IgG (H+L) Highly Cross-Adsorbed Secondary Antibody, Alexa Fluor Plus 647 (Invitrogen, A32795). After the incubation with the secondary antibody, the brain sections were washed with 3X PBS/T followed by 3X 1:1 Ethanol-PBS solution for 5 minutes each. The sections were then incubated in the methoxy-04 solution in dark. The unspecific dye was removed by washing it three times with a 1:1 Ethanol-PBS solution and two times with Milli-Q water, and the slices were then mounted on microscope slides with DAKO fluorescent mounting medium (DAKO, Germany).

### 3D reconstruction of astrocytes and synaptic pruning

The 3D reconstruction of the astrocytes and measurement of synaptic engulfment in the volume of the astrocyte was performed using IMARIS 9.7 (Oxford Instruments, Bitplane) using the protocol as described previously (21). % ROI colocalization measures (proportionally) the amount of synaptic tag engulfed by the entire astrocyte (GFAP) volume. The engulfment analysis was performed to calculate the amount of C1q (eat-me tag) and synaptophysin (pre-synaptic marker) inside the whole volume of individually reconstructed astrocytes. The interaction of plaque with astrocytes was done in a similar method. Instead of individual astrocytes, all the astrocytes in the frame were volumetrically reconstructed and their interaction with the plaques was measured by the percentage of the entire astrocytic volume that interacts with amyloid plaques. The astrocytes that were partially reconstructed or incorrectly rendered were removed manually. On a general note, the astrocyte within a 30 µm radius from the amyloid plaques was considered eligible for the plaque astrocyte interaction. The percentage interaction between C1q and plaque was done using “automatic thresholding” coloc analysis from the IMARIS package.

### Golgi-Cox staining of spines

Animals that went through the chronic or acute treatment for the water cross maze were decapitated using a guillotine and brains were sagitally sectioned to 250 µm hippocampal slices in the vibratome. The slices were recovered for 30min at 35°C in a chamber submerged with artificial cerebrospinal fluid (aCSF) and then at room temperature for an hour. The FD Rapid GolgiStain Kit (FD NeuroTechnologies) was used for Golgi–Cox staining. The staining was performed following the manufacturer’s instructions. The brain slices were incubated in an impregnation solution for four days. Golgi–Cox-stained pyramidal neurons of layer V were imaged for apical dendrites epi-fluorescence microscope (Axio Imager.M2 with ApoTome.2, Jena, Zeiss, Germany). The obtained Z-stacks were opened and analyzed in Fiji (22).

### Steroid profiling by gas chromatography coupled to tandem-mass spectrometry (GC-MS/MS)

A whole panel of steroids was identified and quantified simultaneously in individual tissues by a GC-MS/MS procedure fully validated in terms of accuracy, reproducibility, and linearity in the nervous tissue (23). Briefly, steroids were extracted from the cortex and hippocampus with 10 volumes of MeOH. The following internal standards were added into the extracts for steroid quantification: 2 ng of ^13^C_3_-PROG for PROG, 2 ng of ^13^C_5_-5α-dihydroprogesterone (DHP) (CDN Isotopes, Sainte Foy la Grande, France) for 5α/β-DHP, 2 ng of epietiocholanolone (3β-hydroxy-5β-androstan-17-one, Steraloids, Newport, Rhode Island) for 5α/β-dihydrotestosterone (DHT), 3α/β5α/β-tetrahydrotestosterone (THT), pregnenolone (PREG), 3α/β5α/β-tetrahydroprogesterone (THPROG), 5α/β20α-THPROG, 3α/β5α/β20α/β- hexahydroprogesterone (HHPROG) and 3α/β5α/β-THDOC, (2 ng of 19-norPROG for 5α/β- dihydrodeoxycorticosterone (DHDOC), 10 ng of ^2^H_8_-corticosterone (B) for B and 5α/β- dihydrocorticosterone (DHB), 2 ng of ^13^C_3_-deoxycorticosterone (DOC) for DOC and 1 ng of ^13^C_3_- testosterone(T) for T. 2 ng of ^13^C_5_-20α-DHP (Isosciences) for the analysis of 20α/β-DHP.

Extracts were then purified and fractionated by solid-phase extraction with the recycling procedure by using a C18 cartridge (500 mg, International Sorbent Technology) (Liere et al, 2004). The unconjugated steroids- containing fraction was filtered and fractionated by the HPLC system composed of a WPS-3000SL analytical autosampler, an LPG-3400SD quaternary pump gradient coupled with an SR-3000 fraction collector (Thermoscientific, USA), and a Lichrosorb Diol column (25 cm, 4.6 mm, 5 µm) in a thermostated block at 30 °C. The column was equilibrated in a solvent system of 90% heptane and 10% of a mixture composed of heptane/isopropanol (85/15). Elution was performed at a flow rate of 1 ml/min, first with 90% heptane and 10% of heptane/isopropanol (85/15) for 15 min, then with a linear gradient to 100% acetone in 2 min. This mobile phase was kept constant for 13 min. Two fractions were collected from the HPLC system: 5α/β-DHPROG were eluted in the first HPLC fraction (3-14 min) and were next silylated with 50 μl of a mixture N-methyl-N-trimethylsilyltrifluoroacetamide/ammonium iodide/dithioerythritol (1000:2:5) (vol/wt/wt) for 15 min at 70 °C. The second fraction (14-29 min) containing all other steroids was derivatized with 25 μl heptafluorobutyric anhydride (HFBA) and 25 μl anhydrous acetone for 1 h at 80°C. Both fractions were dried under a stream of N2 and resuspended in heptane.

GC-MS/MS analysis of the purified and derivatized extracts was performed using an AI1310 autosampler, a Trace 1310 gas chromatograph (GC), and a TSQ8000 mass spectrometer (MS) (Thermoscientific, San Jose, CA). The injection was performed in the splitless mode at 280 °C (1 min of splitless time) and the temperature of the gas chromatograph oven was initially maintained at 80 °C for 1 min and ramped between 80 to 200 °C at 20 °C/min, then ramped to 300 °C at 5 °C/min and finally ramped to 350 °C at 30 °C/min. The helium carrier gas flow was maintained constant at 1 ml/min during the analysis. The transfer line and ionization chamber temperatures were 330 °C and 200 °C, respectively. Electron impact ionization was used for mass spectrometry with an ionization energy of 70 eV. Argon was used as the collision gas. GC-MS/MS signals were evaluated using a computer workstation using the software Excalibur®, release 3.0 (Thermoscientific, USA). Identification of steroids was supported by their retention time and according to two or three transitions. Quantification was performed according to the transition giving the more abundant signal. The range of the limit of detection was roughly 0.5-10 pg according to the steroid structure.

The analytical protocol has been validated for all the targeted steroids using extracts of 200 mg of a pool of male mice brains. The evaluation included the limit of detection, linearity, accuracy, intra-assay, and inter-assay precision (23).

### Statistical analysis

All statistical analyses are performed with GraphPad Prism 6 (Version 6.01). The normality of the data sets was checked using the D’Agostino-Pearson and Shapiro–Wilk tests. Where the sample size was smaller, and normality detection was not possible, non-parametric testing was applied. In addition, data sets consisting of less than four samples were not considered for any statistical testing, instead, we use descriptive statistics to note the differences. The data sets are shown with the median and their respective interquartile range except for the escape latency and accuracy curves which are expressed as mean ± SD. The outliers in the sample sets were identified using the ROUT method with Q=1%. For data sets not distributed normally, the significance of differences between experimental groups was accessed by Kruskal–Wallis test followed by Dunn’s multiple comparisons *post hoc* test, and for normally distributed data sets one-way ANOVA followed by Bonferroni’s post-hoc analysis was used. The adjusted *p-value* after the multiple comparisons were calculated and are mentioned in the figure legends. Between two non-normally distributed experimental groups, the differences are analyzed by Mann–Whitney U test. Each figure legend specifies the statistical test used along with the median and interquartile range for each data set. Statistical significance is indicated in the plots with an asterisk (*) when p<0.05.

## Results

### XBD173 prevents the adverse synaptotoxic effects of Aβ_1-42_ on CA1-LTP in the hippocampus via TSPO

LTP can be induced by activating CA1 synapses in the hippocampus briefly with high-frequency electrical stimulation (100 pulses/100 Hz) (24) (**Figure 1 A**). The application of different Aβ oligomers e.g., Aβ_1-42_, 3NTyr10-Aβ, Aβ_1-40,_ and AβpE3 has been shown to impair CA1-LTP in acute hippocampal slices in a concentration-dependent manner (19). Here, we tested whether the application of XBD173 prevents the synaptotoxic effects of Aβ_1-42_ and Aβ_1-40_ oligomers. In our experiments, we applied XBD173 concentration (300nM), which still allows the induction of LTP. We used sagittal hippocampal slices and monitored the fEPSPs in the CA1 stratum radiatum of the hippocampus.

**Figure 1:**
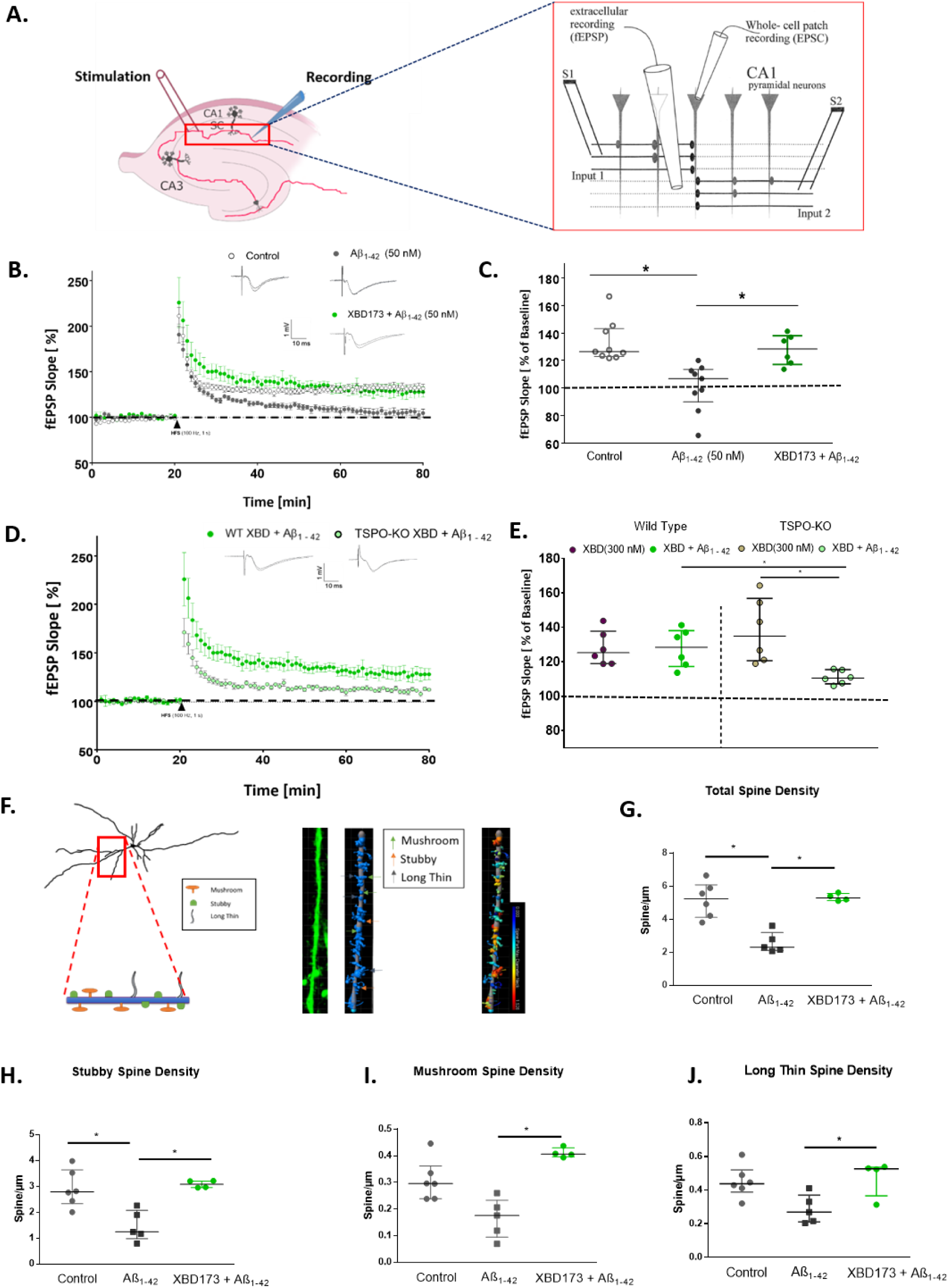
XBD173 via TSPO protein prevents the adverse synaptotoxic effects of Aβ_1-42_ on CA1-LTP in the hippocampus and prevents dendritic spine loss induced by Aβ_1-42_ oligomers. A. Representative schematic for the positioning of electrodes to obtain CA1-LTP in the Schaffer collaterals pathway. The inset referring to the detailed positioning is adapted from (70). B. Normalized field excitatory postsynaptic potential (fEPSP) time course following a high-frequency stimulation (HFS) under control conditions, with 90 min Aβ_1-42_ exposure alone, and the simultaneous application of XBD173 (300 nM) and Aβ_1-42_ (50 nM) respectively. The insets on the top are representative traces for each treatment group. C. Scatter dot plot summarizing the last 10 min (starting from 50 min to 60 min) after HFS for respective groups: Control (n=10), Aβ_1-42_ (n=10), XBD173 + Aβ_1-42_ (n=6) (Kruskal–Wallis test with a Dunn’s multiple comparisons *post hoc* test; Control: 126.5 (123.0-143.3) % of baseline slope vs Aβ_1-42_: 106.9 (89.93-113.6) % of baseline slope, *p=*0.0007; Aβ_1-42_ vs XBD173 + Aβ_1-42_: 128.5 (117.2-138.1) % of baseline slope, *p=*0.0192). D. Normalized field excitatory postsynaptic potential (fEPSP) time course following a high-frequency stimulation (HFS) in WT (XBD173 + Aβ_1-42_), and TSPO-KO (XBD173 + Aβ_1-42_). The insets on the top are representative traces for each treatment group. E. Scatter dot plot summarizing the last 10 min (starting from 50 min to 60 min) after HFS for respective groups: WT XBD173 (n=6), TSPO-KO XBD173 (n=6), WT XBD173 + Aβ_1-42_ (n=6) and TSPO-KO XBD173 + Aβ_1-42_ (n=6) (Mann–Whitney U test: WT XBD173: 125.2 (118.9-137.8) % of baseline slope vs WT XBD173 + Aβ_1-42_:128.5 (117.2-138.1) % of baseline slope, *p=*0.7879; TSPO-KO XBD173: 134.9 (120.6-156.9) % of baseline slope vs TSPO-KO XBD173 + Aβ_1- 42_:110.5 (107.2-115.4) % of baseline slope, *p=*0.0022; WT XBD173 + Aβ_1-42_ vs TSPO-KO XBD173 + Aβ_1-42_, *p=*0.0087). F. XBD173 does not rescue LTP impairment in presence of Aβ_1-42_ in TSPO-KO animals. F. Schematic showing the different spine classifications. At the side of the schematics, the representative apical dendritic segments of the CA1 pyramidal neuron are shown. Dendrites and spines are reconstructed and classified using the Filament function in IMARIS Software. Different classes of spines are indicated with arrows: green: Mushroom, yellow: Stubby, and grey: Long thin spines. The next plot shows a representative categorization of the spines based on the spine neck diameter. G. Total spine density in different treatment groups: Control (n=6), Aβ_1-42_ (n=5), and XBD173 + Aβ_1-42_ (n=4), (Kruskal–Wallis test with a Dunn’s multiple comparisons *post hoc* test; Control: 5.236 (4.108-6.080) spines/ µm vs Aβ_1-42_: 2.302 (2.113-3.194) spines/ µm, *p=*0.0136; Aβ_1-42_ vs XBD173 + Aβ_1-42_: 5.297 (5.134-5.563) spines/ µm, *p=*0.0196) H. Stubby spine density in different treatment groups: Control (n=6), Aβ_1-42_ (n=5), and XBD173 + Aβ_1-42_ (n=4) (Kruskal–Wallis test with a Dunn’s multiple comparisons *post hoc* test; Control: 2.790 (2.332-3.639) spines/ µm vs Aβ_1-42_: 1.244 (0.979-2.078) spines/ µm, *p=*0.0340; Aβ_1-42_ vs XBD173 + Aβ_1-42_: 3.086 (2.956-3.210) spines/ µm, *p=*0.0114). I. Mushroom spine density in different treatment groups: Control (n=6), Aβ_1-42_ (n=5), and XBD173 + Aβ_1-42_ (n=4) (Kruskal–Wallis test with a Dunn’s multiple comparisons *post hoc* test; Control: 0.2959 (0.2390-0.3629) spines/ µm vs Aβ_1-42_: 0.1756 (0.0942-0.2332) spines/ µm, *p=*0.0897; Aβ_1-42_ vs XBD173 + Aβ_1-42_: 0.4063 (0.3966-0.4302) spines/ µm, *p=*0.0049). J. Long-thin spine density in different treatment groups: Control (n=6), Aβ_1-42_ (n=5), and XBD173 + Aβ_1-42_ (n=4) (Kruskal–Wallis test with a Dunn’s multiple comparisons *post hoc* test; Control: 0.4378 (0.3889- 0.5203) spines/ µm vs Aβ_1-42_: 0.2698 (0.2103-0.3712) spines/ µm, *p=*0.0846; Aβ_1-42_ vs XBD173 + Aβ_1-42_: 0.5272 (0.3662-0.5369) spines/ µm, *p=*0.0490). Data are represented as median with their respective interquartile range. *\*p* < 0.05. ns: not significant.

Low nanomolar concentration (50 nM) of both Aβ_1-42_ and Aβ_1-40_ inhibited CA1 LTP after 90 minutes of incubation in the bath solution (**Figure 1 B, C, and Supplementary Figures 1)**. XBD173 (300 nM) preincubation for 60 minutes, prevented LTP impairments induced by Aβ_1-42_ oligomers (**Figure 1 B, C)**. XBD173, however only partly rescues the LTP impairments resulting from the Aβ_1-40_ peptide (**Supplementary Figures 1)**. To access the role of TSPO protein in mediating the neuroprotective effect of XBD173, we also recorded the fEPSPs from the acute hippocampal slices obtained from the global TSPOKO mouse model. XBD173 fails to exert any neuroprotective effect in the transgenic TSPOKO mouse slices incubated with Aβ_1-42_ oligomers indicating a significant involvement of TSPO protein for the beneficial effect of XBD173 on ameliorating Aβ_1-42_-induced synaptic deficits (**Figure 1 D, E)**.

### XBD173 *ex vivo* treatment prevents dendritic spine loss resulting from Aβ_1-42_ oligomers

Dendritic spines are highly dynamic structures and critical for the function of neural circuits (**Figure 1 F)**. An increase in spine density has been reported to accompany LTP and learning and memory-related mechanisms (25). In previous experiments, we showed that Aβ reduces neuronal spines in hippocampal slices, and antagonizing GluN2b receptors reversed this effect (19). Therefore, the application of XBD173 might be a promising approach to antagonize the over-activation of NMDA receptors on pyramidal cells. As such, in line with the beneficial effects of XBD173 against Aβ_1-42_ synaptotoxicity, we intended to unravel whether XBD173 can restore the detrimental effect of Aβ_1-42_ on spine density in hippocampal slices. XBD173 significantly prevented the elimination of spines induced by Aβ_1-42_ oligomers (**Figure 1 G)**. Given the specific functionalities of different classes of spines, we also analyzed the spines according to their classification and quantified in detail stubby, long thin and mushroom spines. XBD173 significantly improved the stubby, mushroom, and long thin spine counts and prevented the synaptotoxic effect of Aβ_1- 42_ oligomers on dendritic spines (**Figure 1 H - J; Supplementary Figure 6)**.

### Chronic but not acute administration of XBD173 ameliorates spatial learning deficits in AD mice via a TSPO-dependent pathway

Given the neuroprotective effects of XBD173 ex vivo, we asked whether XBD173 treatment in transgenic AD-modelled mice may improve cognitive deficits. The WCM was developed as a highly sensitive tool to assess hippocampal-dependent place learning in small animals (20). Water as the only motive force and selective reinforcement paradigm results in more accurate and robust performances than food rewards (20). These properties make this test highly suitable for detecting cognitive deficits developed in ArcAβ mice. The ArcAβ mice model of AD begins to develop robust cognitive impairment at an age group of 6-7 months (26). In this study, we used two different treatment strategies to see whether XBD173 treatment can rescue these cognitive deficits. The treatment schedule of chronic and acute treated animals for the behavioral tests is shown in **Figure 2 A and Supplementary Figure 9 A respectively**. The acute treatment schedule was started 2 days before the start of the training phase in the water cross maze while the chronic treatment of XBD173 (1mg/kg) was followed for twelve weeks from the age of 8 months. In the acute treatment group mice treated with XBD173 had a comparable escape latency as well as accuracy to the vehicle-treated group (**Supplementary Figures 9 A, B)**. Retests 1 and 2 which were accessed a week later and a month later respectively did not show any differences between the XBD173 treatment and the vehicle-treated group (**Supplementary Figures 9 C, D)**. The training phase trajectories for both the escape latency and accuracy were comparable between the experimental groups (**Supplementary Figures 9 A)**. This suggests that acute treatment of XBD173 might not be sufficient to improve spatial learning in the AD model of mice.

**Figure 2:**
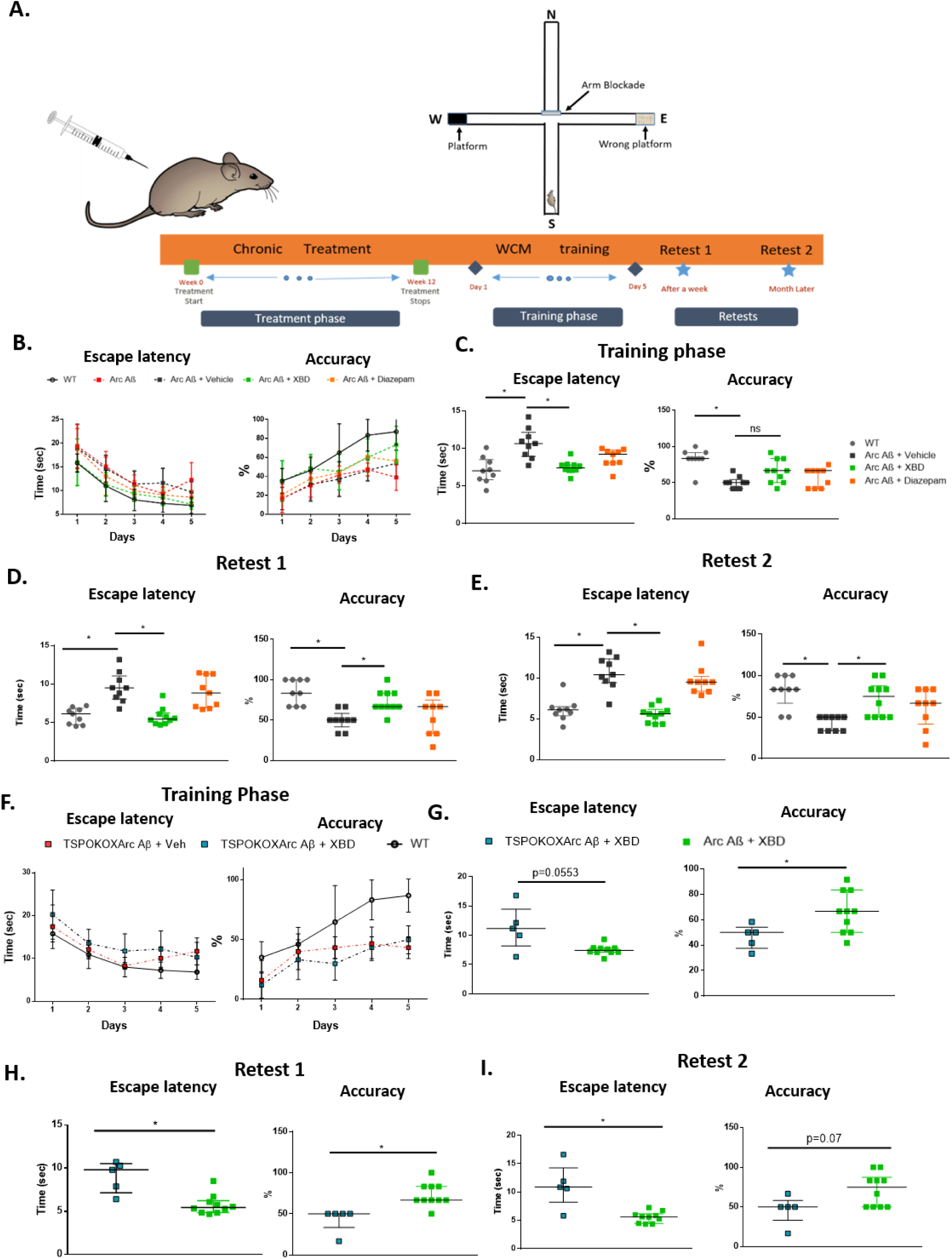
Chronic administration of XBD173 ameliorates cognitive deficits in AD mice via a TSPO- dependent pathway. A. Schematic of chronic treatment schedule, training, and retests in water cross maze. B. The escape latency and accuracy curves during the course of the 5-day training phase in the water cross maze. C. Escape latency and accuracy comparison between the different treatment groups in the training phase. WT (n=9), Arc Aβ + vehicle (n=9), Arc Aβ + XBD (n=10), and Arc Aβ + diazepam (n=9) (Kruskal– Wallis test with a Dunn’s multiple comparisons *post hoc* test; WT: 7 (5.825-8.518) s vs Arc Aβ + vehicle: 10.66 (8.948-12.14) s, *p=*0.0013; Arc Aβ + vehicle vs Arc Aβ + XBD: 7.420 (7.109-7.816) s, *p=*0.0075; Arc Aβ + diazepam: 9.320 (8.073-9.758) s). Accuracy is expressed for each animal as the % of trials correctly performed (Kruskal–Wallis test with a Dunn’s multiple comparisons *post hoc* test; WT: 83.33 (83.33-91.67) % vs Arc Aβ + vehicle: 50 (41.67-54.17) %, *p=*0.0004; Arc Aβ + vehicle vs Arc Aβ + XBD: 66.66 (50-83.33) %, *p=*0.1155; Arc Aβ + diazepam: 66.66 (41.67-66.67) %). D. Escape latency (Kruskal– Wallis test with a Dunn’s multiple comparisons *post hoc* test; WT: 6.113 (4.693-6.829) s vs Arc Aβ + vehicle: 9.477 (8.035-11.05) s, *p=*0.0018; Arc Aβ + vehicle vs Arc Aβ + XBD: 5.430 (4.844-6.237) s, *p=*0.0008; Arc Aβ + diazepam: 8.833 (6.931-11.34) s) and accuracy comparison (Kruskal–Wallis test with a Dunn’s multiple comparisons *post hoc* test; WT: 83.33 (66–100) % vs Arc Aβ + vehicle: 50 (41.67-58.33) %, *p=*0.0003; Arc Aβ + vehicle vs Arc Aβ + XBD: 66.66 (50-66.66) %, *p=*0.0145; Arc Aβ + diazepam: 66.66 (33.33-75) %) between the different treatment groups in the Retest 1 phase. WT (n=9), Arc Aβ + vehicle (n=9), Arc Aβ + XBD (n=10) and Arc Aβ + diazepam (n=9). E. Escape latency (Kruskal–Wallis test with a Dunn’s multiple comparisons *post hoc* test; WT: 6.092 (5.375-6.498) s vs Arc Aβ + vehicle: 10.41 (9.426-12.35) s, *p=*0.0007; Arc Aβ + vehicle vs Arc Aβ + XBD: 5.608 (4.444-6.166) s, *p<*0.0001; Arc Aβ + diazepam: 9.508 (8.363-10.20) s) and accuracy comparison (Kruskal–Wallis test with a Dunn’s multiple comparisons *post hoc* test; WT: 83.33 (66.66-100) % vs Arc Aβ + vehicle: 50 (33.33-50) %, *p=*0.0008; Arc Aβ + vehicle vs Arc Aβ + XBD: 75 (50-87.5) %, *p=*0.0102; Arc Aβ + diazepam: 66.66 (41.67-83.33) %) between the different treatment groups in the Retest 2 phase. WT (n=9), Arc Aβ + vehicle (n=9), Arc Aβ + XBD (n=10) and Arc Aβ + diazepam (n=9). F. The escape latency and accuracy curves during the course of the 5-day training phase in the water cross maze for hetTSPOKO X Arc Aβ + Veh (n=5) and hetTSPOKO X Arc Aβ + XBD (n=5) groups. G. Escape latency (Mann–Whitney U test: hetTSPOKO X Arc Aβ + XBD: 11.16 (8.168-14.46) s vs Arc Aβ + XBD: 7.420 (7.109-7.816) s, *p=*0.0553) and accuracy comparison (Mann–Whitney U test: hetTSPOKO X Arc Aβ + XBD: 50.00 (37.50-54.17) % vs Arc Aβ + XBD: 66.66 (50.00-83.33) %, *p=*0.0296) between the different treatment groups in the training phase. Arc Aβ + XBD (n=10), and hetTSPOKO X Arc Aβ + XBD (n=5) groups. H. Escape latency (Mann– Whitney U test: hetTSPOKO X Arc Aβ + XBD: 9.800 (7.138-10.50) s vs Arc Aβ + XBD: 5.430 (4.844- 6.237) s, *p=*0.0047) and accuracy comparison (Mann–Whitney U test: hetTSPOKO X Arc Aβ + XBD: 50.00 (33.33-50) % vs Arc Aβ + XBD: 66.66 (50-66.66) %, *p=*0.0033) between the different treatment groups in the Retest 1 phase. I. Escape latency (Mann–Whitney U test: hetTSPOKO X Arc Aβ + XBD: 10.86 (8.2-14.24) s vs Arc Aβ + XBD: 5.608 (4.444-6.166) s, *p=*0.008) and accuracy comparison (Mann– Whitney U test: hetTSPOKO X Arc Aβ + XBD: 50.00 (33.33-58.33) % vs Arc Aβ + XBD: 75 (50-87.5) %, *p=*0.07) between the different treatment groups in the Retest 2 phase. Data are represented as median with their respective interquartile range. *\*p* < 0.05. ns: not significant.

We followed up with a chronic treatment schedule to study whether chronic treatment of XBD173 could reverse the spatial learning deficits in the transgenic ArcAβ mice (**Figure 2 A)**. Learning curves and final cognitive performance were markedly poorer in ArcAβ animals than in respective WT as assessed by escape latency and accuracy (**Figure 2 B)**. Learning and accuracy curve trajectory in the training phase shows that chronic treatment of transgenic ArcAβ animals with XBD173 improves spatial learning in AD mice (**Figure 2 B).** Chronic XBD treatment significantly improves the escape latency right away in the training phase (**Figure 2 C)**. While there was a tendency for higher accuracy in the XBD173-treated animals in the training phase, these differences were not significant compared to the vehicle-treated transgenic AD mice (**Figure 2 C)**. Importantly, however, it is essential to note that the cognitive deficits were markedly rescued in both Retest 1 and Retest 2, performed after a week and a month respectively (**Figure 2 D, E)**. The XBD173- treated ArcAβ in both the retests showed significantly better escape latency and accuracy compared to their vehicle-treated transgenic counterparts (**Figure 2 D, E)**. This suggests a long-term neuroprotective effect of XBD173 on cognition even after terminating the treatment for more than a month. Since benzodiazepines are potent anxiolytics and are widely used in clinical practice, we included diazepam as a reference agent and applied this drug in an identical chronical regimen as XBD173. Unlike XBD173, diazepam did not show betterment in the spatial learning paradigm as monitored in the ArcAβ mice group (**Figure 2 D, E)**. The escape latency and accuracy of the diazepam-treated transgenic AD mice were comparable to the vehicle-treated group. Over the training phase and in both the retests, the XBD173-treated group showed a higher percentage of accurate learners compared to the vehicle-treated and diazepam-treated transgenic counterparts (**Supplementary Figures 7 C, D, E)**. It is essential to note that the treatment uniformly affected individual animals, not producing severe deficits in only a few animals as shown in the supplementary figure (**Supplementary Figures 7 A, B)**.

To confirm the involvement of the TSPO protein in the XBD173-mediated neuroprotection in cognition, we crossbred the ArcAβ line with TSPOKO animals. The homozygous TSPOKO X ArcAβ line, however, showed behavioral saliences in the form of epileptic seizures. Therefore, we used the het TSPOKO X ArcAβ to investigate the role of the TSPO protein in conferring cognitive benefits. The het TSPOKO X ArcAβ animals treated with XBD173 did not show any marked improvements in cognitive performance compared to the vehicle-treated group as can be seen from both escape latency and accuracy parameters in the training phase as well as in both the retests (**Supplementary Figures 8)**. The performance of XBD173- treated ArcAβ animals was significantly better than that of het TSPOKO X ArcAβ animals treated either with XBD173 or vehicle (**Figure 2 F - I)**. The latency curve as well as the accuracy trajectory of hetTSPOKO X ArcAβ animals were poor as compared to the WT animals (**Figure 2 F)**. These results indicate the neuroprotective effect of XBD173 on improving spatial learning in the AD mice model is dependent on the TSPO protein.

### Chronic XBD173 administration reduces plaque load, rescues the loss of dendritic spines, and increases neurosteroid levels in AD mice

To investigate the effects of chronic XBD173 treatment on AD pathology we looked at different parameters. We accessed the plaque load after staining with congo derivative methoxy-04 in the hippocampus and cortex of these mice. Interestingly, XBD173-treated mice showed a reduction of plaque load and count particularly in the cortex and not in the hippocampus (**Figure 3 A - E)**. We also quantified soluble Aβ using an Aβ_1-42_-sensitive ELISA. The hemizygous overexpression of APP carrying the ArcAβ mutation increased Aβ_1-42_ levels. Consistent with the plaque load analysis, chronically applied XBD173 changed the amount of soluble Aβ_1-42_ as detected by ELISA only in the cortex (**Figure 3 I, J)**. To understand whether chronic XBD173 treatment rescues the loss of dendritic spines, we analyzed the spine density by Golgi-cox staining. XBD173 treatment significantly rescued the loss of spines in the transgenic (**Figure 3 G, H)**. Further, GC-MS/MS analysis of XBD173-treated brains revealed an increase in several neurosteroid levels such as allopregnanolone (3α,5α-THP), 3β,5α-THDOC and 5α DHDOC in the cortex and hippocampus (**Figure 3 K - N)**. Together, these experiments demonstrate the disease-modifying effect of XBD173 in the AD mouse model. This hypothesis is further supported by reduced plaque load and soluble Aβ_1-42_ levels, rescued synaptic density, and increased neurosteroid levels.

**Figure 3:**
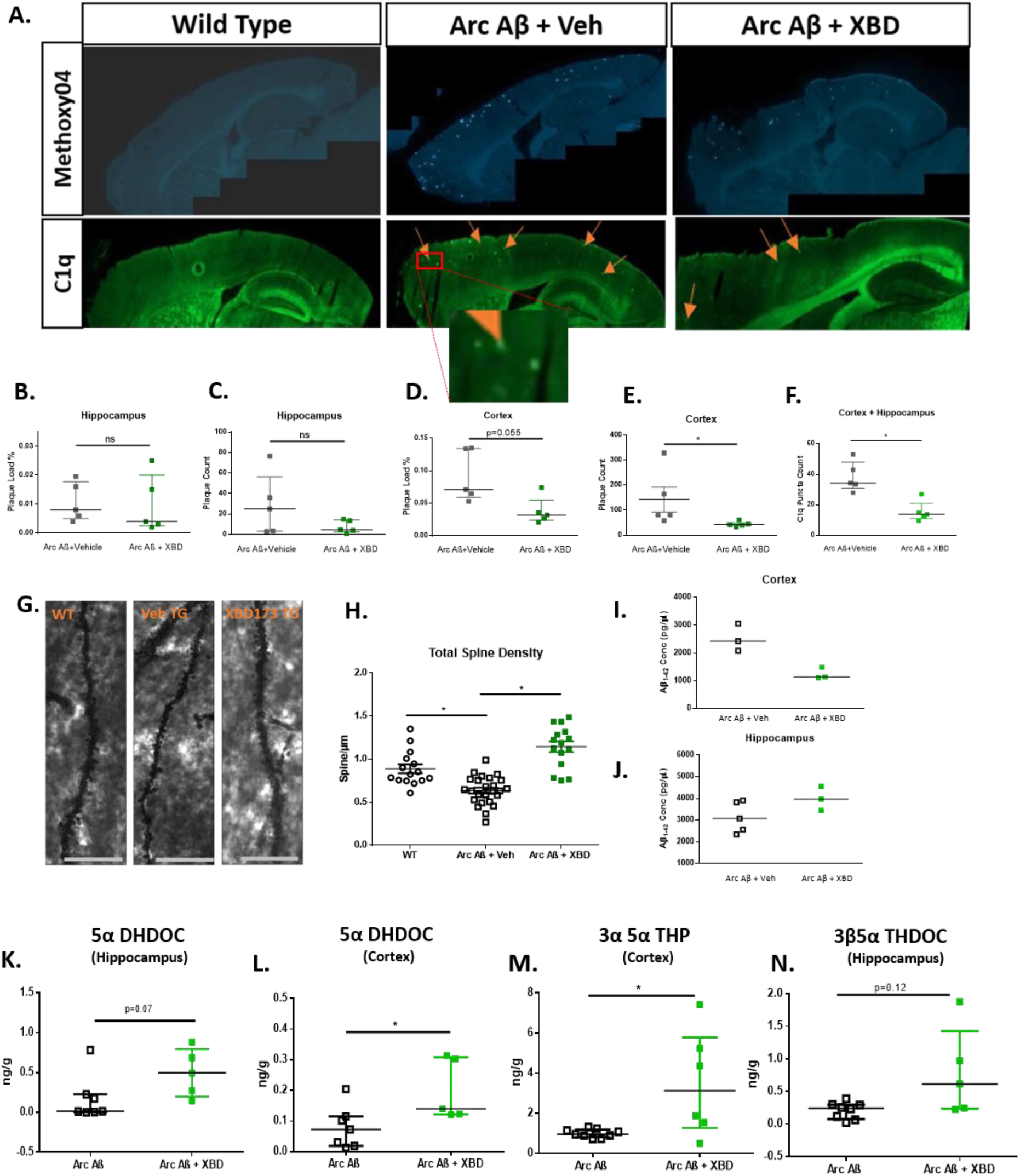
Chronic XBD173 administration reduces plaque load, rescues the loss of dendritic spines, and increases neurosteroid levels in AD mice. A. Representative images for methoxy-04 plaque staining (first row) and C1q puncta staining (second row) in WT, Arc Aβ + vehicle, Arc Aβ + XBD groups. The inset in the second row shows an enlarged view of the C1q puncta. Another 63x magnified image of the puncta colocalized with methoxy-04 plaque is shown in Figure 6. B. Plaque load % comparison between Arc Aβ + vehicle (n=5) and Arc Aβ + XBD (n=5) in the hippocampus (Mann–Whitney U test: Arc Aβ + vehicle: 0.0080 (0.0050-0.01775) % vs Arc Aβ + XBD: 0.0040 (0.0025-0.0200) %, *p=*0.4524). C. Plaque count comparison between Arc Aβ + vehicle (n=5) and Arc Aβ + XBD (n=5) in the hippocampus (Mann–Whitney U test: Arc Aβ + vehicle: 25 (3.25-56.25) vs Arc Aβ + XBD: 4.5 (2.5-14.30), *p=*0.4127). D. Plaque load % comparison between Arc Aβ + vehicle (n=5) and Arc Aβ + XBD (n=5) in the cortex (Mann–Whitney U test: Arc Aβ + vehicle: 0.0710 (0.0590-0.1345) % vs Arc Aβ + XBD: 0.0320 (0.0240-0.0550) %, *p=*0.055). E. Plaque count comparison between Arc Aβ + vehicle (n=5) and Arc Aβ + XBD (n=5) in the cortex (Mann–Whitney U test: Arc Aβ + vehicle: 81.00 (69.50-246.3) vs Arc Aβ + XBD: 41 (36.15-52.50), *p=*0.0159). F. C1q puncta count comparison between Arc Aβ + vehicle (n=5) and Arc Aβ + XBD (n=5) in the cortex and hippocampus (Mann–Whitney U test: Arc Aβ + vehicle: 34.33 (30.75-48.00) vs Arc Aβ + XBD: 14 (11–21), *p=*0.0079). G. Representative Golgi-Cox stained apical dendrites in the hippocampus for WT, Arc Aβ + vehicle, and Arc Aβ + XBD groups respectively. H. Total spine density calculated from n=15-25 dendrites obtained from three to four mice per group (one-way ANOVA with a Bonferroni’s multiple comparisons *post hoc* test; WT: 0.8244 (0.7545-1.008) spines/ µm vs Arc Aβ + vehicle: 0.6329 (0.5177-0.7629) spines/ µm, *p=*0.0005; Arc Aβ + vehicle vs Arc Aβ + XBD: 1.190 (0.9424-1.320) spines/ µm, *p*<0.0001). I. Soluble Aβ_1-42_ levels were measured in the cortex. WT (n=3), Arc Aβ + vehicle (n=3), and Arc Aβ + XBD (n=3). J. Soluble Aβ_1-42_ levels measured in the hippocampus WT (n=3), Arc Aβ + vehicle (n=5), and Arc Aβ + XBD (n=3). K. 5α DHDOC levels in the hippocampus for the treatment groups from GC-MS/MS analysis Arc Aβ + vehicle (n=7) and Arc Aβ + XBD (n=5) (Mann–Whitney U test: Arc Aβ + vehicle: 0.0150 (0.0060-0.2270) ng/g vs Arc Aβ + XBD: 0.4940 (0.2115-0.7860) ng/g, *p=*0.07). L. 5α DHDOC levels in the cortex for the treatment groups from GC-MS/MS analysis Arc Aβ + vehicle (n=7) and Arc Aβ + XBD (n=5) (Mann–Whitney U test: Arc Aβ + vehicle: 0.0730 (0.0190-0.1150) ng/g vs Arc Aβ + XBD: 0.1400 (0.1220-0.3085) ng/g, *p=*0.0177). M. 3α 5α THP levels in the cortex for the treatment groups from GC-MS/MS analysis Arc Aβ + vehicle (n=9) and Arc Aβ + XBD (n=6) (Mann–Whitney U test: Arc Aβ + vehicle: 0.9490 (0.8125-1.188) ng/g vs Arc Aβ + XBD: 3.127 (1.273-5.791) ng/g, *p=*0.036). N. 3β5*α* THDOC levels in the hippocampus for the treatment groups Arc Aβ + vehicle (n=8) and Arc Aβ + XBD (n=5) (Mann–Whitney U test: Arc Aβ + vehicle: 0.2415 (0.076-0.2955) ng/g vs Arc Aβ + XBD: 0.6130 (0.2350-1.427) ng/g, *p=*0.12). Data are represented as median with their respective interquartile range. *\*p* < 0.05. ns: not significant.

### Ex vivo XBD173 treatment does not affect the microglial polarization states which are altered via Aβ_1-42_ oligomers

Previous studies have demonstrated a shift in microglial polarization (an increase in M1 microglia and a reduction in M2 microglia) in AD (27, 28). We therefore asked whether this shift in the polarization states occurs after the addition of Aβ_1-42_ oligomers in the slice and whether ex vivo XBD173 incubation could potentially stop this shift. While in general there was a tendency for an increase in protein expression as measured by the mean fluorescence intensity (MFI) of different microglial markers (CX3CR1 and TMEM119) after treatment with Aβ_1-42_ oligomers, a significant increase in MFI was only visible for the microglia marker P2RY12 (**Supplementary Figures 5 C - E)**. The microglial polarization states were quantified by respective M1 (CD80) or M2 (CD163) markers. We observed an increased expression of both M1 and M2 microglial polarization markers after subsequent treatment with Aβ_1-42_ oligomers. However, XBD173 treatment did not alter the elevated CD80 or CD163 levels (**Supplementary Figures 5 A, B)**. These findings suggest that ex vivo XBD173 treatment does not affect microglial polarization.

### Chronic XBD173 treatment reduces aberrant astrocytic phagocytosis and pruning of synapses

While TSPO is expressed in both neuronal and glial cells, the upregulation of astrocytic TSPO occurs prior to the microglial TSPO upregulation in AD pathology (16). Therefore, astrocytes can be considered an essential target for TSPO ligands. Astrocytes are known to actively eliminate synapses via synaptic pruning in developing and maturing neurons. This pruning process, however, is exacerbated in the case of AD pathophysiology.As such, we hypothesized whether chronic XBD173 treatment could reduce the increased astrocytic pruning in transgenic AD mice. When analyzing high-resolution individual astrocytes, we observed that chronic XBD173 treatment significantly reduced the enhanced synaptic engulfment by astrocytes in both the cortex and hippocampus **(Figure 4)**. This suggests a lower removal of functional synapses contributing to better cognitive function in XBD173-treated mice.

**Figure 4:**
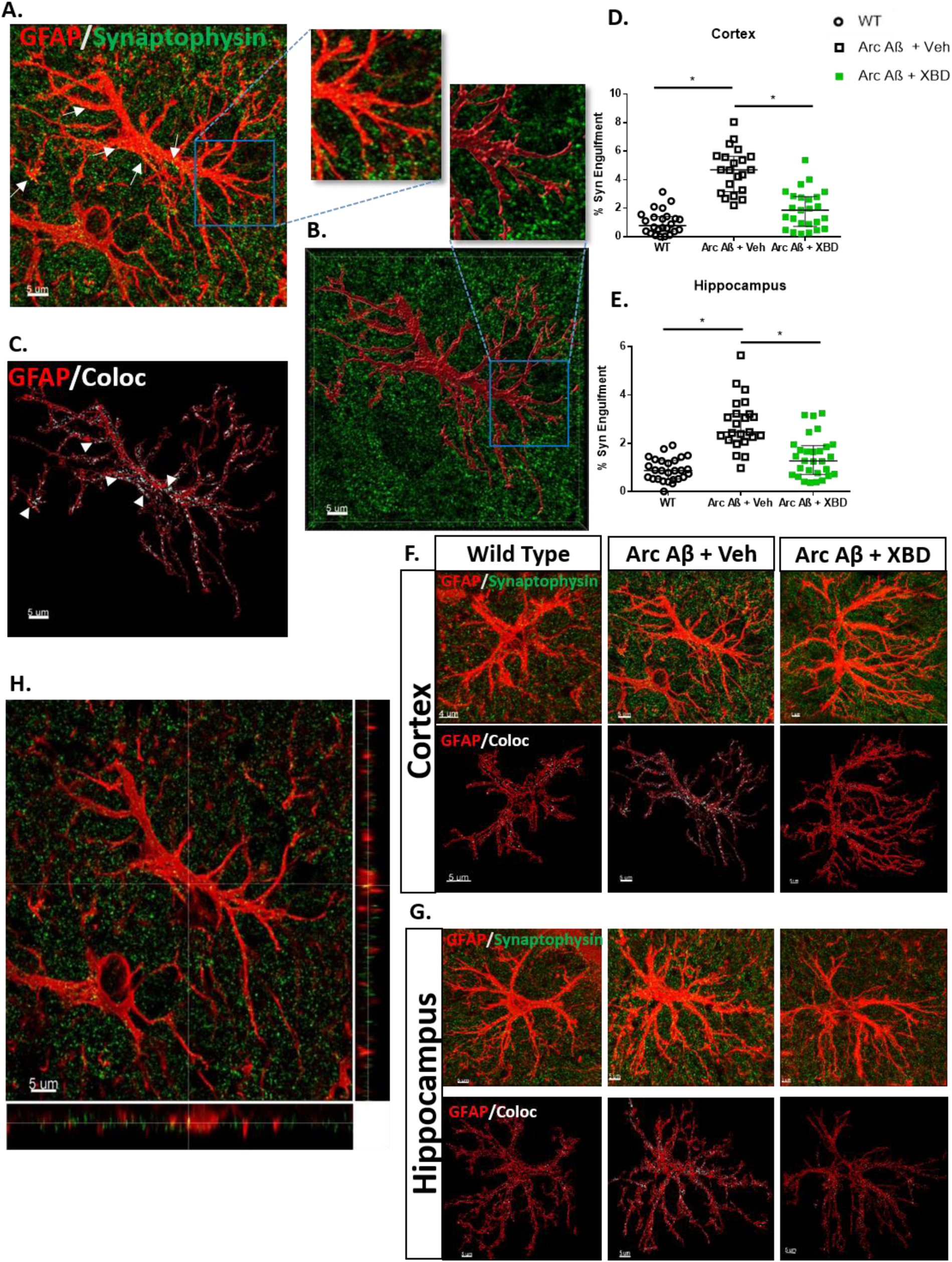
Chronic XBD173 treatment reduces aberrant astrocytic phagocytosis and pruning of synapses. A. Individual high-resolution astrocytes (shown in red) and pre-synaptic marker synaptophysin (shown in green). The colocalization points are indicated by an arrow. B. Volumetric reconstruction of astrocytes (red) using IMARIS 9.7 with synaptophysin (green). The insets show enlarged sections of before and after reconstruction of the astrocytes. C. Rendered outline of astrocyte (red) with colocalization between astrocytes and synaptophysin marked in white and shown by the arrow. D. % of Synaptophysin engulfment by astrocyte quantified in different treatment groups in the cortex (one-way ANOVA with a Bonferroni’s multiple comparisons *post hoc* test; WT: 0.7809 (0.3763-1.411) % vs Arc Aβ + vehicle: 4.692 (3.157- 5.640) %, *p<*0.0001; Arc Aβ + vehicle vs Arc Aβ + XBD: 1.866 (0.7195-2.818) spines/ µm, *p*<0.0001). E. % of Synaptophysin engulfment by astrocyte quantified in different treatment groups in the hippocampus (Kruskal–Wallis test with a Dunn’s multiple comparisons *post hoc* test; WT: 0.8785 (0.5336-1.293) % vs Arc Aβ + vehicle: 2.468 (2.144-3.232) %, *p<*0.0001; Arc Aβ + vehicle vs Arc Aβ + XBD: 1.276 (0.7177- 1.906) %, *p*<0.0001; 21-29 astrocytes collected from 4-5 mice per group). F. Representative images for GFAP (red) and Synaptophysin (green) in the cortex in different experimental groups. The first panel shows the original image and the second panel shows the colocalization point (white) inside the rendered volume of the astrocyte from the respective groups. G. Representative images for GFAP (red) and Synaptophysin (green) in the hippocampus in different experimental groups. The first panel shows the original image and the second panel shows the colocalization point (white) inside the rendered volume of the astrocyte from the respective groups. H. Orthogonal plane showing the GFAP (red) and synaptophysin (green). *Scale bar: 5 µm*. Data are represented as median with their respective interquartile range. *\*p* < 0.05. ns: not significant.

### Chronic XBD173 treatment decreases the C1q puncta in AD mice and reduces the aberrant increase in astrocytic engulfment of C1q complement protein

In physiological conditions, several studies have reported a C1q-independent astrocytic pruning of synapses (29). However, in pathological conditions such as AD, there is strong evidence for C1q-dependent removal of synapses by the astrocytes (29). To further understand the effects of XBD173 on astrocytic pruning in AD, we studied the connection of C1q with the astrocytic engulfment of synapses. We found deposition of C1q in form of a puncta-like structure which highly colocalized with amyloid plaques in the transgenic ArcAβ mice (**Figure 5 E)**. However, while all the C1q puncta were associated with the amyloid plaques, not all the plaques were associated with these puncta. We asked whether chronic XBD173 treatment affects the colocalization percentage of C1q puncta with the plaques. Colocalization of C1q puncta with the plaques was comparable between both XBD173 and vehicle-treated groups (**Figure 5 F)**. However, XBD173- treated group showed a significantly lower number of C1q puncta as compared to their vehicle-treated counterparts (**Figure 2 F)**. Reduction in these C1q puncta in XBD173-treated animals, could possibly explain the reduction of aberrant increase of complement protein C1q in AD transgenic mice.

**Figure 5:**
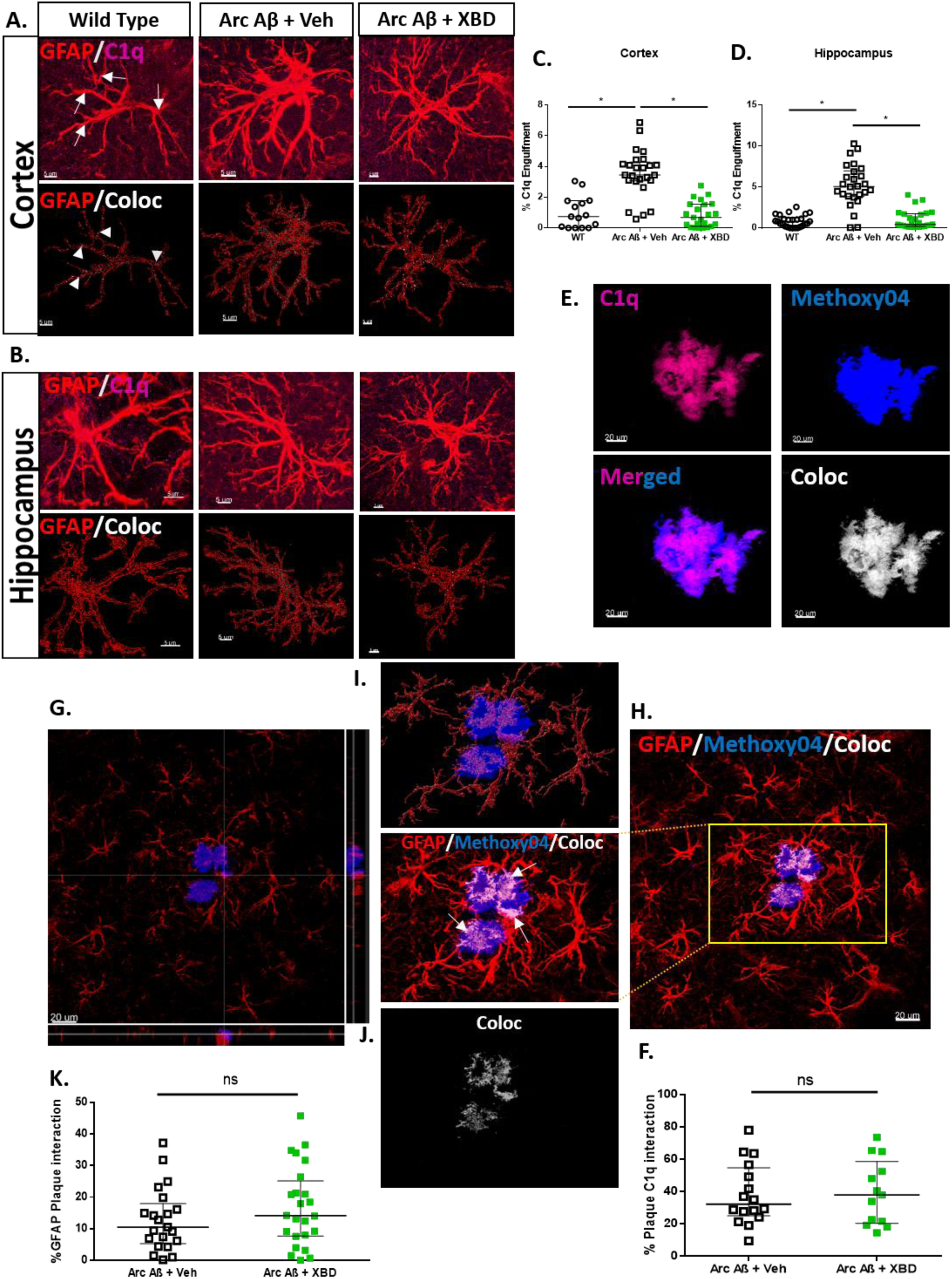
Chronic XBD173 treatment reduces the aberrant increase in astrocytic engulfment of C1q complement protein. A. Representative images for GFAP (red) and C1q (magenta) in the cortex in different experimental groups. The first panel shows the original image and the second panel shows the colocalization point (white) inside the rendered volume of the astrocyte from the respective groups. *Scale bar: 5 µm.* B. Representative images for GFAP (red) and C1q (magenta) in the hippocampus in different experimental groups. The first panel shows the original image and the second panel shows the colocalization point (white) inside the rendered volume of the astrocyte from the respective groups. *Scale bar: 5 µm .* C. % of C1q engulfment by astrocyte quantified in different treatment groups in the cortex (Kruskal–Wallis test with a Dunn’s multiple comparisons *post hoc* test; WT: 0.7465 (0.0033-1.767) % vs Arc Aβ + vehicle: 3.448 (3.071-4.300) %, *p<*0.0001; Arc Aβ + vehicle vs Arc Aβ + XBD: 0.6829 (0.1004- 1.543) %, *p*<0.0001; 15-23 astrocytes collected from 4-5 mice per group). D. % of C1q engulfment by astrocyte quantified in different treatment groups in the hippocampus (Kruskal–Wallis test with a Dunn’s multiple comparisons *post hoc* test; WT: 0.8102 (0.1865-1.250) % vs Arc Aβ + vehicle: 5.056 (3.849- 6.986) %, *p<*0.0001; Arc Aβ + vehicle vs Arc Aβ + XBD: 0.4372 (0.1979-1.733) %, *p*<0.0001; 26 astrocytes per collected from 5 mice per group). E. Representative 63X images of C1q puncta (magenta), methoxy-04 plaques (blue), merged, and colocalization (white) in transgenic AD mice. *Scale bar: 20 um.*F. % plaque C1q interaction in different treatment groups. G. Orthogonal plane representation of GFAP (red) and Plaque (blue) interaction (Mann–Whitney U test: Arc Aβ + vehicle: 32.18 (25.07-54.76) % vs Arc Aβ + XBD: 37.99 (20.39-58.68) %, *p=*0.9390; 13-16 astrocytes collected from 4 mice per group). H. Representative image for GFAP (red) and Plaque (blue) interaction. The inset shows an enlarged section of the interaction with colocalization marked by white arrows. I. Reconstructed astrocytes (red) after removal of background signal along with methoxy-04 (blue). J. The colocalization of GFAP and methoxy-04 plaque is shown in white. *Scale bar: 20 µm.* K. % plaque GFAP interaction in different treatment groups in the cortex and hippocampus. Arc Aβ + vehicle and Arc Aβ + XBD (Mann–Whitney U test: Arc Aβ + vehicle: 10.50 (5.305-17.99) % vs Arc Aβ + XBD: 14.15 (7.715-25.15) %, *p=*0.3573; analysis from n=21-24 plaques obtained from 4 mice per group). respective groups. H. Orthogonal plane showing the GFAP (red) and synaptophysin (green). *Scale bar: 5 µm*. Data are represented as median with their respective interquartile range. *\*p* < 0.05. ns: not significant.

C1q has been implicated as an “eat-me tag” for the neurons to undergo elimination. Since XBD173 treatment reduced the aberrant elimination of synapses by astrocytes, we hypothesized that individual astrocytes in the transgenic mice would subsequently engulf more of this C1q “eat-me tag” signatures and XBD173 treatment could affect this astrocytic engulfment of C1q. Consistent with this hypothesis, we observed increased engulfment of C1q by astrocytes in the vehicle-treated transgenic group. XBD173 treatment significantly reduced the astrocytic engulfment of C1q protein in the cortex and hippocampus (**Figure 6 A - D; Supplementary Figures 10)**. Taken together, these results highlight C1q puncta as an additional biomarker for AD and show that XBD173 treatment could rescue the exacerbated C1q capture by astrocytes.

**Figure 6:**
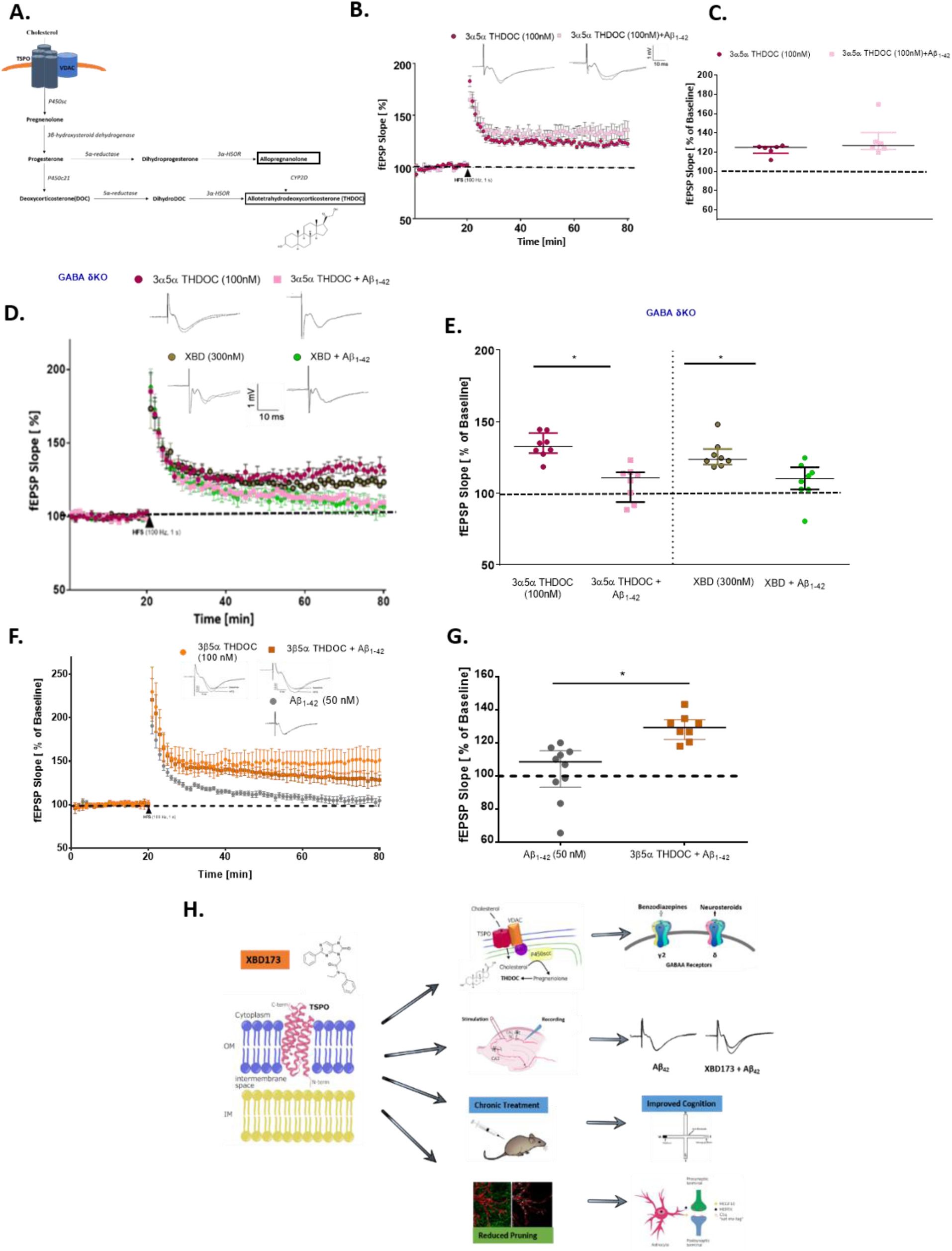
XBD173 promotes a TSPO-mediated synthesis of THDOC and other neurosteroids which enhances GABA_A_ receptor activity containing GABA delta subunit A. Representative schematic of neurosteroid synthesis regulated by the TSPO protein. B. Normalized field excitatory postsynaptic potential (fEPSP) time course following a high-frequency stimulation (HFS) under THDOC (100 nM) and THDOC + Aβ_1-42_ conditions in WT C57/Bl6 mice. The insets on the top are representative traces for each treatment group. C. Scatter dot plot summarizing the last 10 min (starting from 50 min to 60 min) after HFS for respective groups: THDOC (n=6) and THDOC + Aβ_1-42_ (n=6) (Mann–Whitney U test: WT THDOC: 125.0 (119.0-126.0) % of baseline slope vs WT THDOC + Aβ_1-42_: 127.1 (122.9-140.5) % of baseline slope, *p=*0.3874) D. Normalized field excitatory postsynaptic potential (fEPSP) time course following a high- frequency stimulation (HFS) under THDOC (100 nM), THDOC + Aβ_1-42_, XBD173 (300 nM) and XBD173 + Aβ_1-42_ conditions in transgenic GABA-δ KO mice. E. Scatter dot plot summarizing the last 10 min (starting from 50 min to 60 min) after HFS for respective groups in GABA-δ KO mice: THDOC (100 nM) (n=7), THDOC + Aβ_1-42_ (n=7), XBD173 (300 nM) (n=6) and XBD173 + Aβ_1-42_ (n=6) (THDOC: 132.7 (128.1-142.1) % of baseline slope vs THDOC + Aβ_1-42_: 110.8 (93.8-114.7) % of baseline slope, *p=*0.0003; XBD: 123.7 (120.1-130.9) % of baseline slope vs XBD + Aβ_1-42_: 110.2 (102.7-118.2) % of baseline slope, *p=*0.0070). F. Normalized field excitatory postsynaptic potential (fEPSP) time course following a high- frequency stimulation (HFS) under 3β5α THDOC (100 nM), Aβ_1-42_ (50 nM) and 3β5α THDOC + Aβ_1-42_. G. Scatter dot plot summarizing the last 10 min (starting from 50 min to 60 min) after HFS for respective groups in WT C57/Bl6 mice: Aβ_1-42_ (50 nM) (n=10) and 3β5α THDOC + Aβ_1-42_ (n=8) (Mann-Whitney U test; Aβ_1-42_: 108.5 (93.18-115.3) % of baseline slope vs 3β5α THDOC + Aβ_1-42_: 129.3 (122.1-134) % of baseline slope, *p<*0.0001). H. Proposed working mechanism for TSPO-dependent XBD173. Data are represented as median with their respective interquartile range. *\*p* < 0.05. ns: not significant.

### Astrocytic phagocytosis does not contribute to the clearance of methoxy plaques in the cortex

In view of the decreased plaque load in the cortex of XBD173-treated animals and its effect on astrocytic phagocytosis, we asked whether astrocytic phagocytosis of plaques may result in reduced plaque load in these animals. We investigated the number of plaques inside the surrounding astrocytic volume. Astrocytic density around the plaques (at a radius of 30 µm from the plaques) was higher than more away from the plaques. We found that the astrocytic engulfment of plaque was comparable for both vehicle and XBD173- treated animals (**Figure 5 G - K)**. These results indicate that astrocytic phagocytosis does not directly contribute to the removal/reduced plaque load in XBD173-treated transgenic AD mice.

### The TSPO ligand XBD173 promotes the synthesis of different neurosteroids which, increases GABA_A_ receptor activity containing the GABA_A_ receptor delta subunit and provide neuroprotection against Aβ oligomers

Previously TSPO ligands have been attributed to increasing neurosteroidogenesis (30) (**Figure 6 A)**. Given the importance of TSPO dependency of XBD173 as well as the increase in neurosteroids from the chronic treatment of XBD173, we investigated the role of different neurosteroids in preventing LTP impairment. TSPO ligands were previously shown to elevate the levels of the neurosteroids allopregnanolone and pregnenolone (31–33). We started with different concentrations of neurosteroids and monitored the fEPSPs response to see whether these neurosteroids could mimic the neuroprotective action of XBD173. Both pregnenolone (100 nM) and allopregnanolone (10 nM and 30 nM), were not able to prevent the LTP impairments from Aβ_1-42_ oligomers (**Supplementary Figures 2, 3)**. However, allopregnanolone (100 nM) was able to prevent the LTP impairment resulting from the Aβ_1-40_ peptide (**Supplementary Figures 3)**. In addition, 3α5α THDOC (100 nM) was able to rescue the LTP impairment resulting from Aβ_1-42_ oligomers (**Figure 6 B, C)**. Although both neurosteroids and benzodiazepines are positive allosteric modulators of GABA_A_ receptors they bind to different subunits. For instance, while benzodiazepines are sensitive to γ2 subunits, neurosteroids are sensitive to the δ subunit (34). To test our hypothesis on neurosteroid dependence of XBD173-mediated effects on ameliorating the Aβ_1-42_ mediated synaptotoxicity, we used acute hippocampus slices from transgenic animals containing GABA-δ KO. 3α5α THDOC (100 nM) was not able anymore to exert a beneficial action on the LTP in these slices. Additionally. XBD173 (300 nM) failed to exert the neuroprotective effect on GABA-δ KO hippocampal slices after treatment with the Aβ_1-42_ oligomers (**Figure 6 D, E)**. These data suggest that the GABA_A_ receptor containing the δ subunit is required for the neuroprotective effects off the TSPO ligand XBD173. The chronic treatment of XBD173 also results in increase of 3β5α THDOC which is a steroisomer of 3α5α THDOC. We therefore also checked whether 3β5α THDOC isomer was able to rescue the LTP impairment. Similar to its 3α stereoisomer, 3β5α THDOC also prevented the LTP impairment resulting from the incubation of Aβ_1-42_ oligomers (**Figure 6 F, G)**. All these evidence suggests that XBD173 orchestrates the formation of several neurosteroids which play a crucial role in providing comprehensive neuroprotection against the Aβ derieved pathophysiology.

## Discussion

TSPO ligands are neuroprotective in a range of neurological and psychiatric disorders (13, 35). One promising candidate is the TSPO ligand XBD173 (AC-5216/emapunil) which exerts rapid anxiolytic effects in animal models and humans (17). In this study, we provide both in vitro and in vivo evidence for the neuroprotective effect of XBD173 in AD via the TSPO protein. In our study, we show that XBD173 confers neuroprotection against Aβ_1-42_ oligomers and partly rescues the LTP impairment occurring from Aβ_1-40_ oligomers. This differential action of XBD173 could be attributed to the mechanistic differences in how both Aβ_1-42_ and Aβ_1-40_ act to inhibit the formation of CA1 LTP in the hippocampus. In addition, XBD173 rescued the loss of mushroom spines as well as the long thin spines. While mushroom spines because of a higher head volume are thought to play a critical role in long-term memory-related functions, long thin spines because of their adaptability are also called learning spines (36). This suggests that the synaptic dysfunction of neurotransmission resulting from the Aβ_1-42_ oligomer is prevented by the administration of XBD173.

The water cross maze provides conclusive evidence that XBD173 betters spatial learning in the transgenic ArcAβ mice and is dependent on the TSPO to mediate this beneficial action. This, however, challenges the recent reports from Shi et al., which suggest benzodiazepines and TSPO ligands alter synaptic plasticity and cause cognitive impairment (37). Respective differences could be due to the differences in experimental conditions used in both the studies. First, pathological expression of TSPO is different from the physiological state (38, 39). As described in the discussion section later, XBD173 could downtune the noise from tonically overactive glutamatergic signal in AD via GABA_A_ containing delta subunit, thereby leading to a better plasticity/learning profile. In a physiological condition as studied by Shi et al., (37), however, this could lead to the imbalance of the glutamatergic and GABAergic systems, leading to impaired plasticity. This falls in line with our observation of control chronic treatment groups where the wild-type mice chronically treated with XBD173 have lower accuracy in Retest 2 (not significant) as compared to the untreated counterparts (Data not shown). Second, the dose of XBD173 (5 mg/kg) used by Shi et al., (37), as well as the duration (consecutive days), varies considerably from our study (1mg/kg every alternate day). It is therefore essential that both hypo- and hyper-activation of the glutamatergic system leads to neuronal dysfunction and there has to be a subtle balance in any therapeutics that are designed for AD (40).

The finding of the beneficial role of XBD173 has clinical relevance, particularly in peri-operative anesthesia. Benzodiazepines owing to their fast-acting and effective anxiolytic/sedative characteristics are extensively used in perioperative anesthesia (41). These, however, have been associated with undesired side-effect profile, which includes tolerance development, anterograde amnesia, withdrawal symptoms as well as addiction (37,42–45). The administration of benzodiazepine midazolam has been associated with increasing the clinical progression of AD by impacting the tau phosphorylation in a rodent model of AD (46). In animal models and humans, TSPO ligand XBD173 has been shown to exert rapid anxiolytic effects without sedation, tolerance development, and withdrawal symptoms (17). XBD173 can cross the blood-brain barrier (47). Additionally, it has been shown to be neuroprotective in a rodent model of retinal ischemia (48). Considering the role of TSPO in neurosteroidogenesis and the positive allosteric modulation of GABA_A_ receptors by neurosteroids, XBD173 might constitute an alternative to benzodiazepines in perioperative anesthesia.

Chronic administration of XBD in the ArcAβ decreased the plaque load, particularly in the cortex, and also reduced the soluble Aβ levels in the cortex. This effect was not seen in the hippocampus. The transgenic ArcAβ mice did not form extensive plaques in the hippocampus region even at age of around 12 months old. The TSPO expression pattern is not homogeneous and differs across the brain regions, adding to it the cellular expression of TSPO in a pathologic state, which could be the reason for regional differences. A decrease in plaque load and plaque count in the cortex of XBD-treated animals indicates either an increased clearance of the plaques or protection machinery which avoids plaque formation. Chronic XBD173 treatment also rescued spine loss and thereby preserved functional spines.

Glial cells, astrocytes, and microglia play a crucial role in maintaining synaptic homeostasis (49). Loss of functional synapses due to injury or disease affects neuronal communication and leads to network disruption in the central nervous system (CNS). Reactive astrogliosis has often been characterized as a hallmark feature of AD (50). Importantly, TSPO upregulation in astrocytes occurs prior to that in other cells, making astrocytes an interesting target to look at while designing therapeutic strategies. Moreover, activation of the complement C3 may be crucial for the pathophysiology of AD via increased synapse elimination both in human AD as well as a mouse model of tauopathy (51–53). In previous studies, TSPO ligands have been shown to reduce reactive gliosis in mouse models of neurodegeneration (54). Consistent with this, in our model the TSPO ligand XBD173 reduced the phagocytic engulfment of presynaptic particles which was significantly upregulated in the transgenic mouse. The reduction in phagocytic engulfment was consistent in both the hippocampus and the cortex. Previous reports show impairment of the astrocytic phagocyte receptors Multiple EGF Like Domains 10 (MEGF10) and Mer Tyrosine Kinase (MERTK) in the murine AD model, which implies an inefficient clearance of senile plaques (55). We, therefore, asked whether the reduction in the number of amyloid plaques in the cortex region of XBD173- treated mice was due to better phagocytic clearance of these amyloid plaques. Interestingly, we observed that TSPO activation does not significantly change the GFAP-stained astrocytes-plaque interaction, which suggests a similar astrocytic recruitment profiling around the plaque. Moreover, the uptake of the plaque by the astrocyte is not altered by the treatment with XBD173. This suggests that TSPO activation by XBD173 does not affect the phagocytic clearance of the senile plaque, but instead confers neuroprotection by interfering with the plaque formation machinery and stopping the formation of senile plaques. A second possible way would be the activation of microglia either directly by XBD173 or indirectly via astrocytes to clear up the amyloid plaques.

The complement system together with glial cells is crucial for mediating early synapse loss thereby driving the pathophysiology of AD (56–58). In the developing neurons, C1q and C3 of the complement pathway localize to synapses and facilitate the refining of synapses by pruning (59). This process however becomes aberrant in AD pathology with several reports suggesting an abnormal increase in C1q levels of the murine model of AD (56). MEGF10, which is expressed on the astrocytic surface, is a receptor for C1q, which in turn attaches to the phosphatidylserine expressed on the apoptotic cell surface (60). Previously, C1q deletion has been associated with a marked reduction in astrocyte-synapse association, thereby rescuing the synaptic density (29). Similarly, injection of oligomeric Aβ has been shown to increase the association of C1q and postsynaptic markers (56, 61). C1q has also been reported to have a specific binding site for Aβ (62). Both C1q and oligomeric Aβ act in an overlapping way to eliminate the synapses and treatment of the hippocampal slices with anti-C1q antibody significantly rescues the impairment in the LTP caused by the oligomeric Aβ (56, 63). All these studies point towards C1q playing a crucial role as a destruction signal protein in synapse elimination. For these reasons, we studied whether the activation of TSPO protein by XBD173 and its effect on reduced astrocytic phagocytosis is related to the astrocytic engulfment of C1q “eat-me tags” in the AD model. In the current study, we found that the astrocyte engulfment of C1q is enhanced in the hippocampus and the cortex of the AD mouse model. This possibly explains the increase in the loss of functional synapses and an impairment of cognitive function that is evident from the behavioral tests. Chronic treatment of XBD173 decreased this increased engulfment of C1q “eat me tags” in the hippocampus. The reduction in astrocytic engulfment of both presynaptic marker (synaptophysin) and eat-me tag (C1q) suggests that the XBD173 treatment directly reduces the abnormal loss of functional synapses. Taken together, the results from our study as well as previous reports support the hypothesis that C1q is an interesting target for developing effective therapeutics in AD. Targeting of C1q not only affects the proinflammatory status of glial cells but also affects the cross-communication between both astrocytes and microglia.

We also noted the deposition of C1q in form of distinct puncta heavily in the cortex of the transgenic AD mice. These C1q puncta highly overlap with the Aβ plaques (52). While the report of colocalization of amyloid plaques with C1q of the complement pathway remains scarce, one of the previous studies shows distinct colocalization of Aβ protein with C3 complement protein in certain substructural domains within drusen in the retina (64). Few of the previous reports have also shown a colocalization of both C4 binding protein as well as factor H of the complement pathway with the amyloid plaques and dead cells in the AD brain (65). Therefore, the puncta formation could be explained by the deposition of dystrophic neurites. It is interesting to note that not all the plaques overlap with these C1q puncta but all the C1q puncta do overlap with amyloid plaques. This might be of relevance because first, this might act as an additional diagnostic marker to amyloid plaque, and second, this could give us an overview/role of complement-mediated inflammation in the pathophysiology of AD. Taken together, the complement protein C1q may play play a critical role in pathogenesis of AD.

TSPO ligands such as etifoxine have shown neuroprotective effects in multiple sclerosis models via an increase in allopregnanolone levels (66). Similarly, levels of neurosteroids such as allopregnanolone and pregnenolone have been shown to be associated with the expression of the TSPO protein. Consistent with the role of TSPO in the neurosteroidogenesis, we observed elevated levels of 3α5α THP in the cortex, 3ꞵ5α THDOC in the hippocampus, and 5α DHDOC levels in the cortex and hippocampus of the chronic XBD- treated animals. This suggests that XBD173 might affect a multitude of neurosteroids via TSPO and these neurosteroids target a multitude of pathophysiological features of AD. We, therefore, studied the neuroprotective effect of neurosteroids in preventing Aβ_1-42_ mediated LTP deficit. Allopregnanolone prevents LTP impairment resulting from the Aβ_1-40_ oligomer. On the other hand, XBD173 was able to partly rescue the LTP impairment from Aβ_1-40_ oligomers. This could be explained by a shorter duration/concentration of XBD173 in an *ex vivo* setup which might not produce desirable protection as 100 nM allopregnanolone is applied directly to the bath solution. Moreover, certain 3β neurosteroids are known for their functional antagonism of 3α-reduced neurosteroids at GABA_A_ receptors. (67). We found that 3β,5α-THDOC, which was elevated after chronic treatment of XBD173, also prevents the LTP disruption resulting from Aβ_1-42_ oligomers. Additionally, its stereoisomer 3α,5α-THDOC (100 nM) similar to XBD173 prevented the LTP impairment from the Aβ_1-42_ oligomer.

The extrasynaptic benzodiazepine-insensitive GABA_A_ receptors containing the δ-subunits are considered important targets for neurosteroids. In our study, we found that knocking out the δ-subunit of the GABA_A_ receptor impairs the neuroprotective effect of 3α,5α-THDOC against the Aβ_1-42_ oligomer. Importantly, XBD173 similar to THDOC could not prevent the LTP impairment from Aβ oligomer in the absence of δ- subunit containing GABA_A_ receptors. These results provide two essential insights. 1. TSPO-dependent XBD173 acts via the GABA_A_ receptor containing the δ-subunit to exert its neuroprotective effects. 2. The downstream activity of XBD173 is possibly mediated by neurosteroids such as THDOC which acts via the δ-subunit. As such, Aβ has opposing effects on the NMDA receptors in contrast to APP i.e. Aβ increases the glutamate concentration in the synaptic cleft and has an agonistic effect on the NMDA receptors (40). Contrary to the physiological state where the NMDA receptors need strong depolarization of the postsynaptic membrane by glutamate (mM conc.) for the removal of voltage-dependent Mg^2+^ ion blockade, in AD, NMDA receptors are activated for longer durations by a much lower concentration of glutamate (µM conc.) (40). LTP or synaptic plasticity largely depends on the detection of relevant signals over the background noise from the moderate intracellular Ca^2+^ signal (higher signal-to-noise ratio). This signal-to-noise ratio however is impaired in presence of the Aβ oligomers due to increased noise from the chronically overactive glutamatergic system and impaired Mg^2+^ filter in the NMDA receptors (40). Previously, it has been reported that extrasynaptic GABA_A_ receptors are well suited to antagonize the over-activation of NMDA receptors on pyramidal cells (68). Taken together, one could speculate that XBD173 modulation of the δ subunit GABA_A_ receptor could down tune this background noise from the overactivated glutamatergic system in presence of Aβ oligomers. This would therefore also relieve the energy burden due to increased neuronal activity in the presence of Aβ oligomers.

In conclusion, the present study suggests the neuroprotective properties of XBD173 in a mouse model of AD. Nevertheless, some limitations should be considered. First, while the ArcAβ mice model recapitulates the major phenotypes of AD including plaque formation, cognitive impairment, or reactive gliosis, it lacks the presence of neurofibrillary tangles. Second, one could perceive the anxiolytic effect generated by XBD173 administration as a primary mediator of improvement in the cognitive deficit in the water cross maze. While it is difficult to completely separate the neuroprotective from the anxiolytic effects, the performance of XBD173-treated animals in two different retests tested months apart even after the termination of the treatment points towards the beneficial effects on cognitive performance. Moreover, it is essential to note that XBD173 improves the LTP deficit in the ex vivo hippocampal slices which is a cellular correlate of the learning and memory function. In summary, the present results indicate the beneficial effects of XBD173 against Aβ-derived pathology (**Figure 6 H)**. Chronic XBD173 treatment ameliorates cognition and suggests a disease-modifying effect when applied at the early stages of AD, which may be due to. XBD173-induced plaque reduction, soluble Aβ levels, and reduced synaptic pruning.

## Acknowledgment

The work was supported by German Research Foundation (Deutsche Forschungsgemeinschaft) (DFG) research grant (RA 689/12-1) to GR and DFG project number 422179811 to RR. We would like to acknowledge the scientific illustrations obtained from the publically available domain (Scidraw) (69). We would like to thank Dr. Ritu Mishra (Core Facility, Translatum) at Klinikum Rechts der Isar for access to the confocal microscopy facility and the Imaris analysis system. We would like to thank Dr. Carsten Wotjak for his suggestions in WCM experiment.

## Supplementary Figures

**Supplementary Figure 1:**
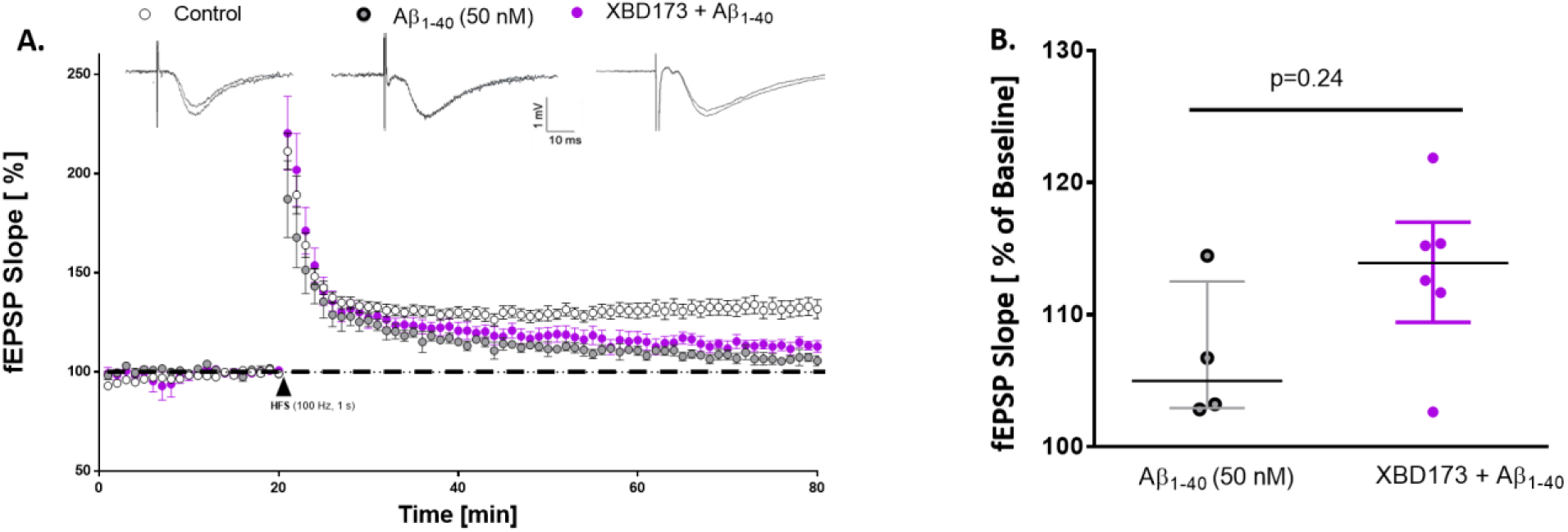
XBD173 partly rescues the LTP impairments resulting from Aβ_1-40_ oligomers. A. Normalized field excitatory postsynaptic potential (fEPSP) time course following a high-frequency stimulation (HFS) under control conditions, with 90 min Aβ_1-40_ exposure alone, and the simultaneous application of XBD173 (300 nM) and Aβ_1-40_ respectively. The insets on the top are representative traces for each treatment group. C. Scatter dot plot summarizing the last 10 min (starting from 50 min to 60 min) after HFS for respective groups: Control (n=10), Aβ_1-40_ (n=4) and XBD173 + Aβ_1-40_ (n=6)(Mann-Whitney U test; Aβ_1-40_: 105 (102.9-112.5) % of baseline slope vs XBD173 + Aβ_1-40_: 113.9 (109.4-117) % of baseline slope, *p=*0.24). Data are represented as median with their respective interquartile range. *\*p* < 0.05. ns: not significant.

**Supplementary Figure 2:**
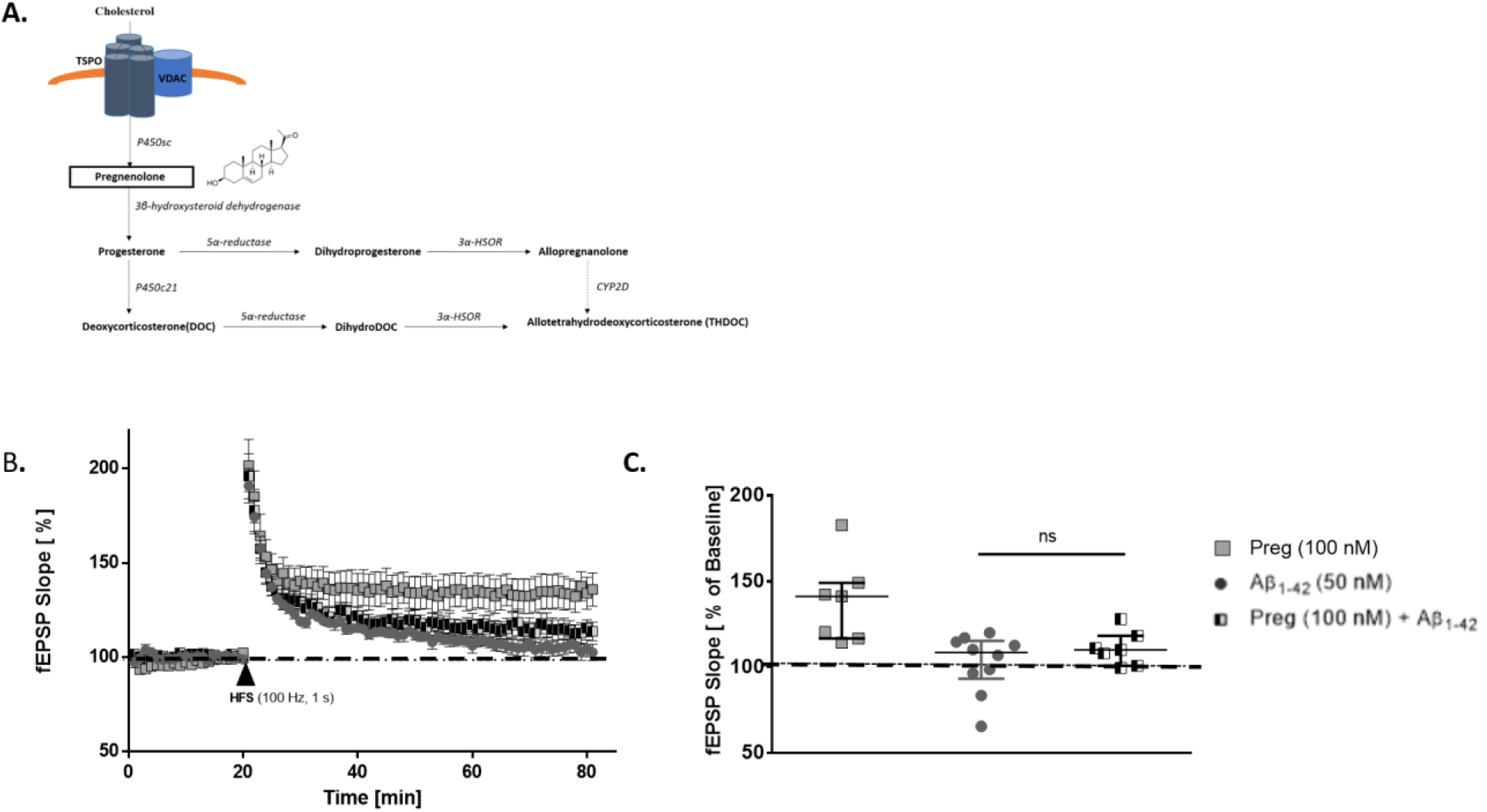
Pregnenolone does not rescue LTP impairments induced by Aβ_1-42_ oligomers. A. Representative schematic of neurosteroid synthesis regulated by TSPO B. Normalized field excitatory postsynaptic potential (fEPSP) time course following a high-frequency stimulation (HFS) under pregnenolone (Preg) (100 nM), Aβ_1-42_ (50 nM) and Preg+ Aβ_1-42_. C. Scatter dot plot summarizing the last 10 min (starting from 50 min to 60 min) after HFS for respective groups in WT C57/Bl6 mice: Preg (100 nM) (n=7), Aβ_1-42_ (50 nM) (n=10) and Preg+ Aβ_1-42_ (n=7) (Mann-Whitney U test; Aβ_1-42_: 108.5 (93.18- 115.3) % of baseline slope vs Preg (100 nM) + Aβ_1-42_: 110 (100.8-118.3) % of baseline slope, *p=*0.47). Data are represented as median with their respective interquartile range. *\*p* < 0.05. ns: not significant.

**Supplementary Figure 3:**
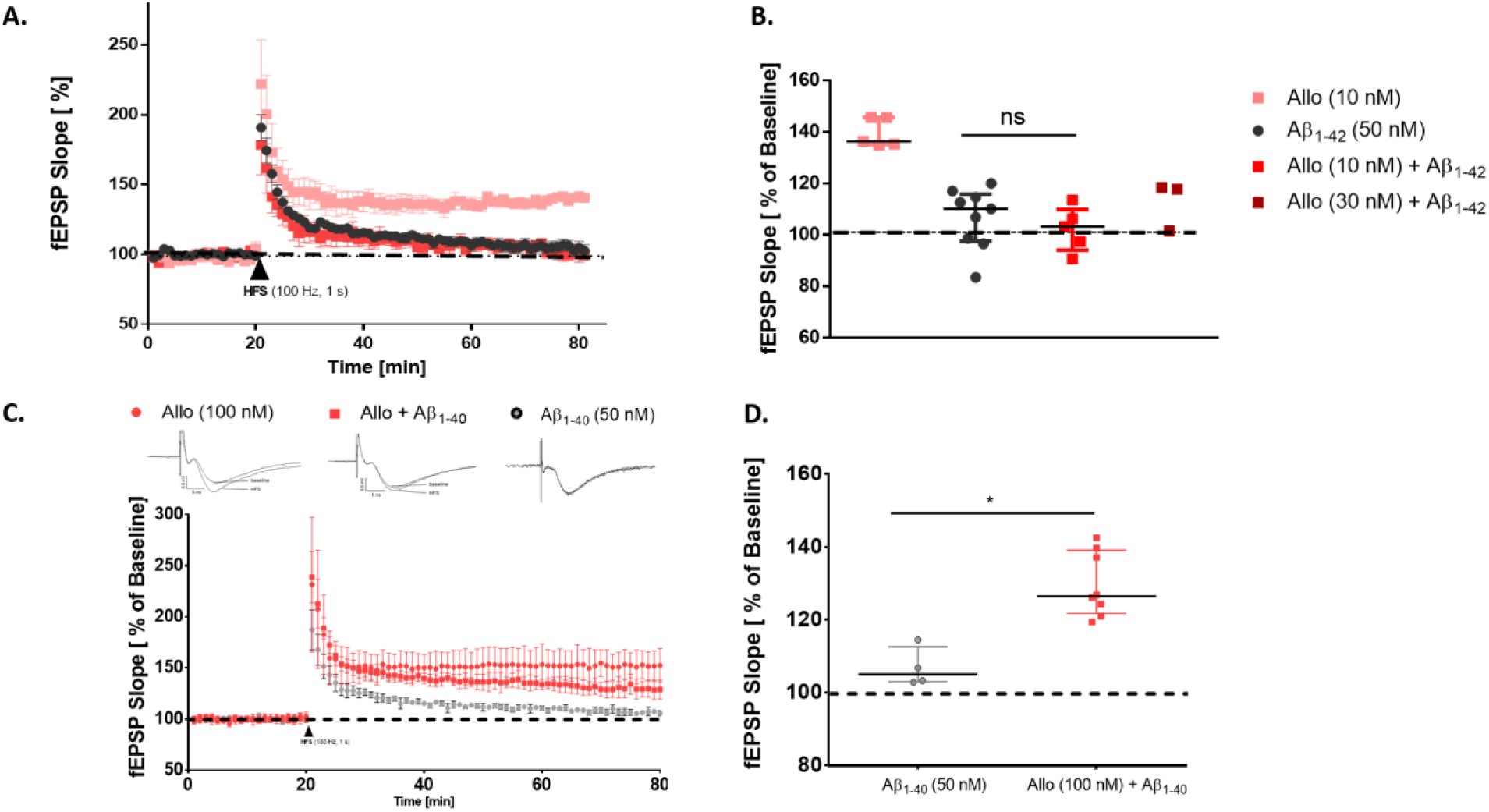
Allopregnanolone (Allo) doesn’t rescue the LTP impairments resulting from Aβ_1-42_ oligomers but protects against the LTP impairment from Aβ_1-40_ oligomers. A. Normalized field excitatory postsynaptic potential (fEPSP) time course following a high frequency stimulation (HFS) under Allo (10 nM), Aβ_1-42_ (50 nM), Allo (10 nM)+ Aβ_1-42_, and Allo (30 nM)+ Aβ_1-42_ . B. Scatter dot plot summarizing the last 10 min (starting from 50 min to 60 min) after HFS for respective groups in WT C57/Bl6 mice: Allo (10 nM) (n=5), Aβ_1-42_ (50 nM) (n=9), Allo (10 nM)+ Aβ_1-42_ (n=5), and Allo (30 nM)+ Aβ_1-42_ (n=3) (Mann-Whitney U test; Aβ_1-42_ : 110.2 (97.55-115.9) % of baseline slope vs Allo (10 nM) + Aβ_1-42_ : 103.2 (94.08-109.9) % of baseline slope, *p=*0.34). C. Normalized field excitatory postsynaptic potential (fEPSP) time course following a high frequency stimulation (HFS) under Allo (100 nM), Aβ_1-40_ (50 nM) and Allo + Aβ_1-40_ . D. Scatter dot plot summarizing the last 10 min (starting from 50 min to 60 min) after HFS for respective groups in WT C57/Bl6 mice: Aβ_1-40_ (50 nM) (n=4) and Allo (100 nM) + Aβ_1- 40_ (n=8) (Mann-Whitney U test; Aβ_1-40_ : 105 (102.9-112.5) % of baseline slope vs Allo (100 nM) + Aβ_1-40_: 126.4 (121.8-139.1) % of baseline slope, *p=*0.004). Data are represented as median with their respective interquartile range. *\*p* < 0.05. ns: not significant.

**Supplementary Figure 4:**
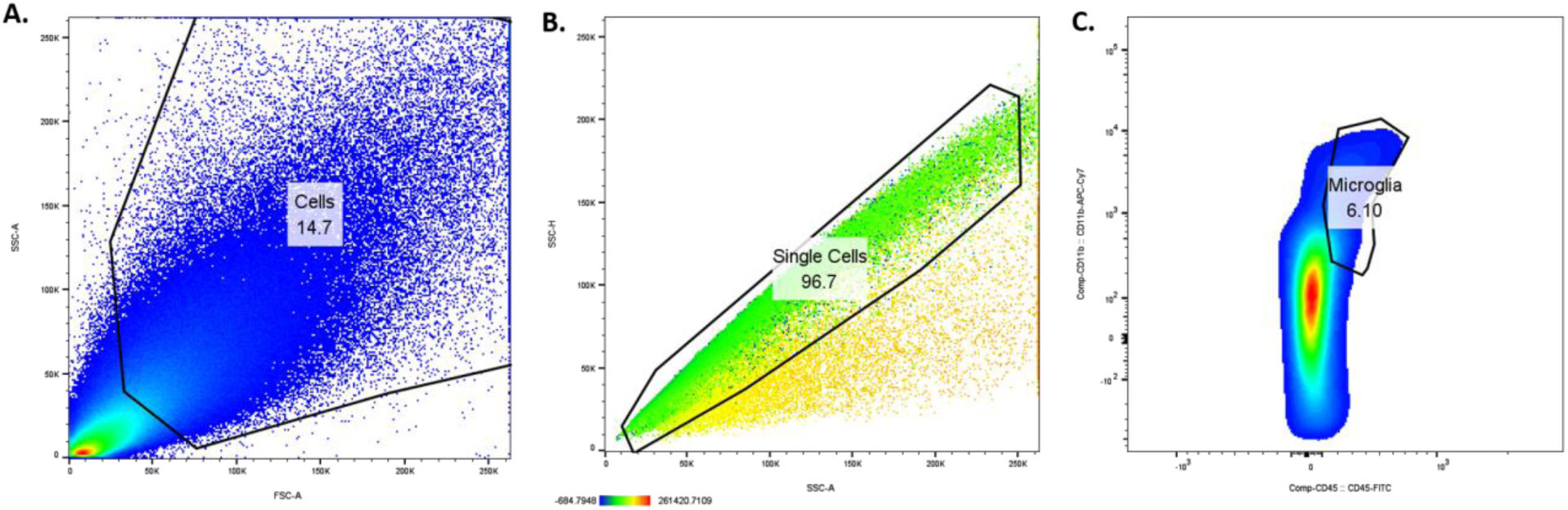
FACS gating strategy to study microglial polarization. A) Cell gating: Front scatter (FCS-A) vs. side scatter (SSC-A) dot plot B) Single-cell gating: SSC-H vs. SSC-A dot plot C) Microglial gating: CD45-FITC vs. CD11b-APC-Cy7 fluorescence intensity dot plot: smoothened, gate for CD45low-to-intermediate and CD11b+ cells

**Supplementary Figure 5:**
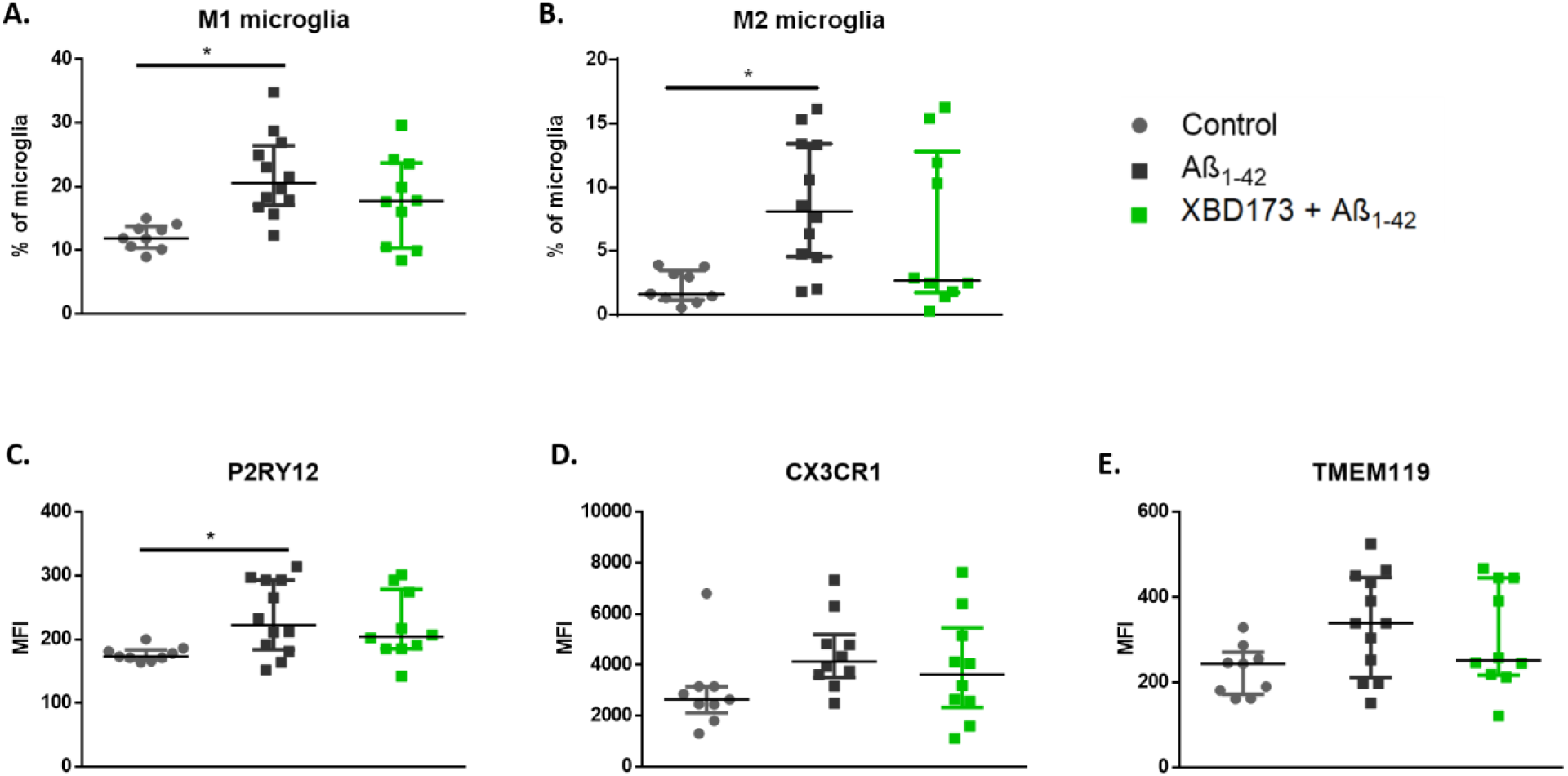
Ex vivo XBD173 treatment does not affect the microglial polarization states altered by Aβ_1-42_ oligomers. A. % of M1 microglia in the cortex and hippocampus in the different treatment groups. Control (n=9), Aβ_1-42_ (n=12) and XBD173 + Aβ_1-42_ (n=10) (Kruskal–Wallis test with a Dunn’s multiple comparisons *post hoc* test; Control: 11.9 (10.35-13.75) % vs Aβ_1-42_: 20.55 (17.08-26.4) %, *p=*0.0013; Aβ_1-42_ vs XBD173 + Aβ_1-42_: 17.70 (10.35-23.68) %, *p=*0.3081). B. % of M2 microglia in the cortex and hippocampus in different treatment groups. Control (n=9), Aβ_1-42_ (n=12) and XBD173 + Aβ_1-42_ (n=10) (Kruskal–Wallis test with a Dunn’s multiple comparisons *post hoc* test; Control: 1.660 (1.155-3.495) % vs Aβ_1-42_: 8.110 (4.55-13.38) %, *p=*0.0068; Aβ_1-42_ vs XBD173 + Aβ_1-42_: 2.705 (1.740-12.78) %, *p=*0.3662). Mean fluorescence intensities (MFI) from FACS measurement for P2RY12 (Kruskal–Wallis test with a Dunn’s multiple comparisons *post hoc* test; Control: 173 (168.5-183.5) vs Aβ_1-42_: 222.5 (183.8- 293), *p=*0.0162; Aβ_1-42_ vs XBD173 + Aβ_1-42_: 204.5 (185-278.8), p>0.99). (C), CX3CR1 (D), and TMEM119 (E) for different treatment groups. Data are represented as median with their respective interquartile range. *\*p* < 0.05. ns: not significant.

**Supplementary Figure 6:**
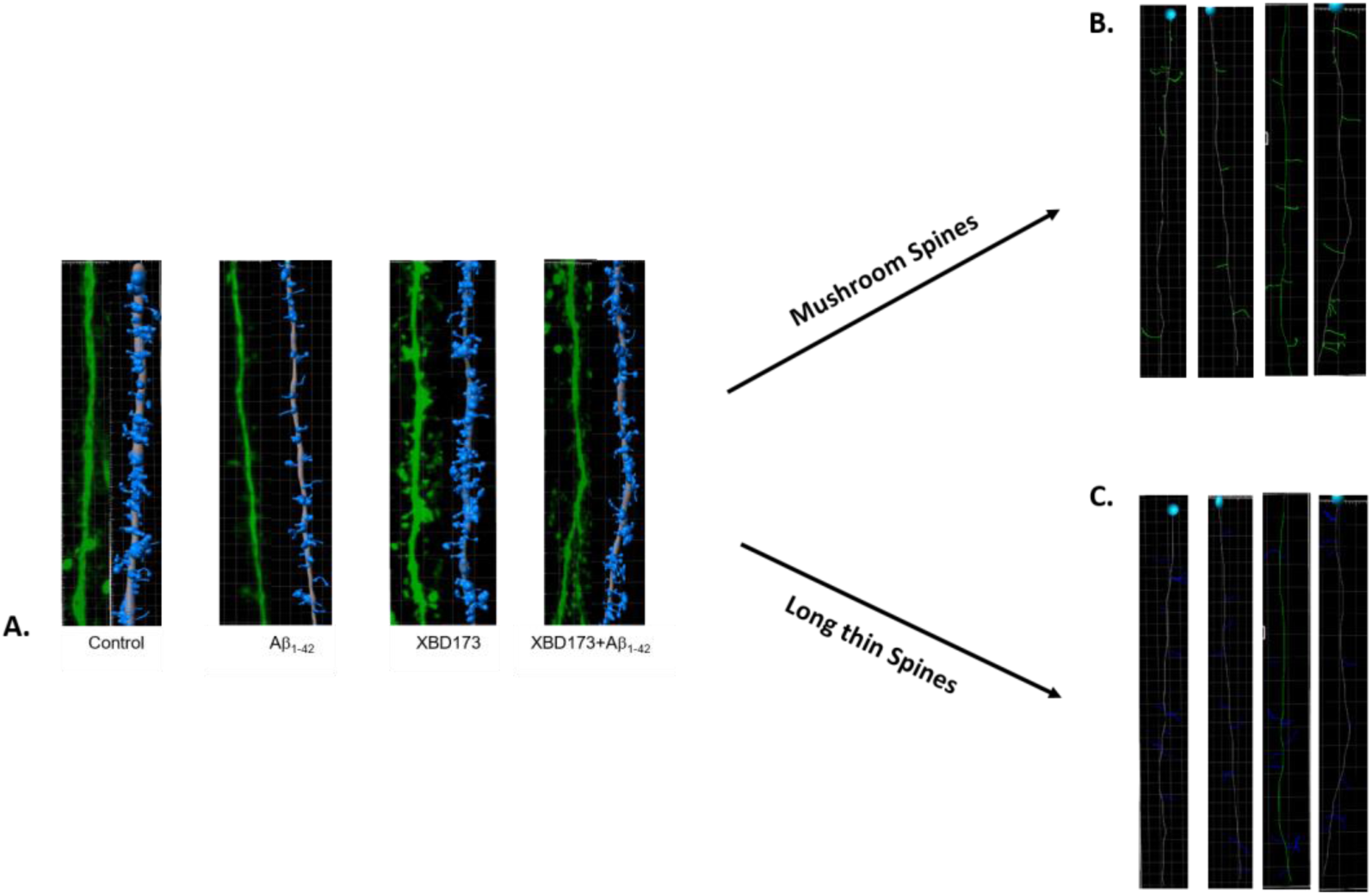
A. Representative images of dendritic spines along with its rendered image from IMARIS. Control (n=6), Aβ_1-42_ (n=5), XBD173 (n=4), and XBD173 + Aβ_1-42_ (n=4) B. After categorization into Mushroom spines for different treatment groups. C. After categorization into long thin spines for different treatment groups.

**Supplementary Figure 7:**
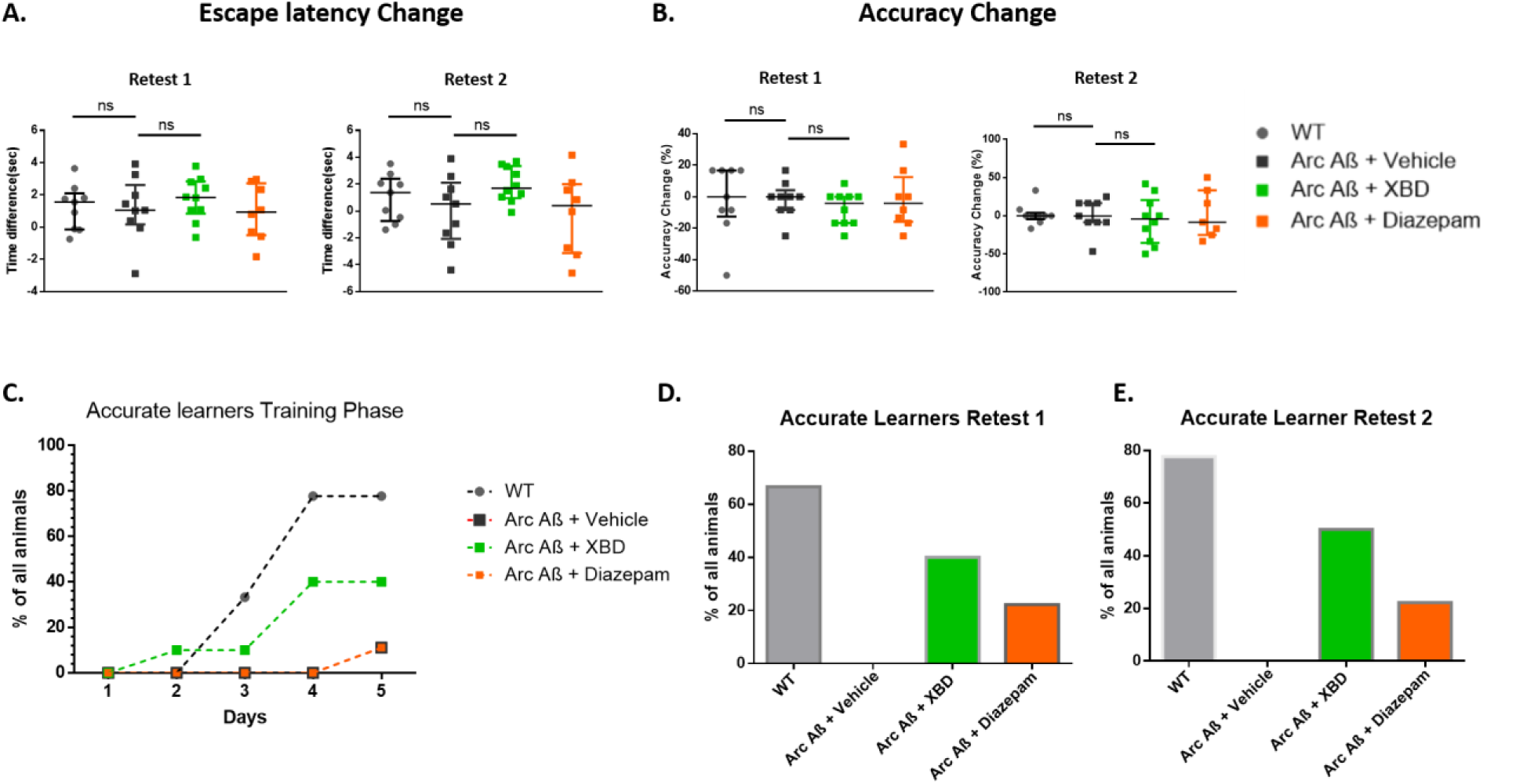
A. Escape latency change in Retest 1 and 2 from the training phase for the different chronic treatment groups. WT (n=9), Arc Aβ + vehicle (n=9), Arc Aβ + XBD (n=10) and Arc Aβ + diazepam (n=8). B. Accuracy change in Retest 1 and 2 from the training phase for the different chronic treatment groups. WT (n=9), Arc Aβ + vehicle (n=9), Arc Aβ + XBD (n=10) and Arc Aβ + diazepam (n=8). C. Accurate learners during the 5-day training phase for different treatment groups. D. Bar plot showing accurate learners in Retest 1. E. Bar plot showing accurate learners in Retest 2. Data are represented as median with their respective interquartile range. *\*p* < 0.05. ns: not significant.

**Supplementary Figure 8:**
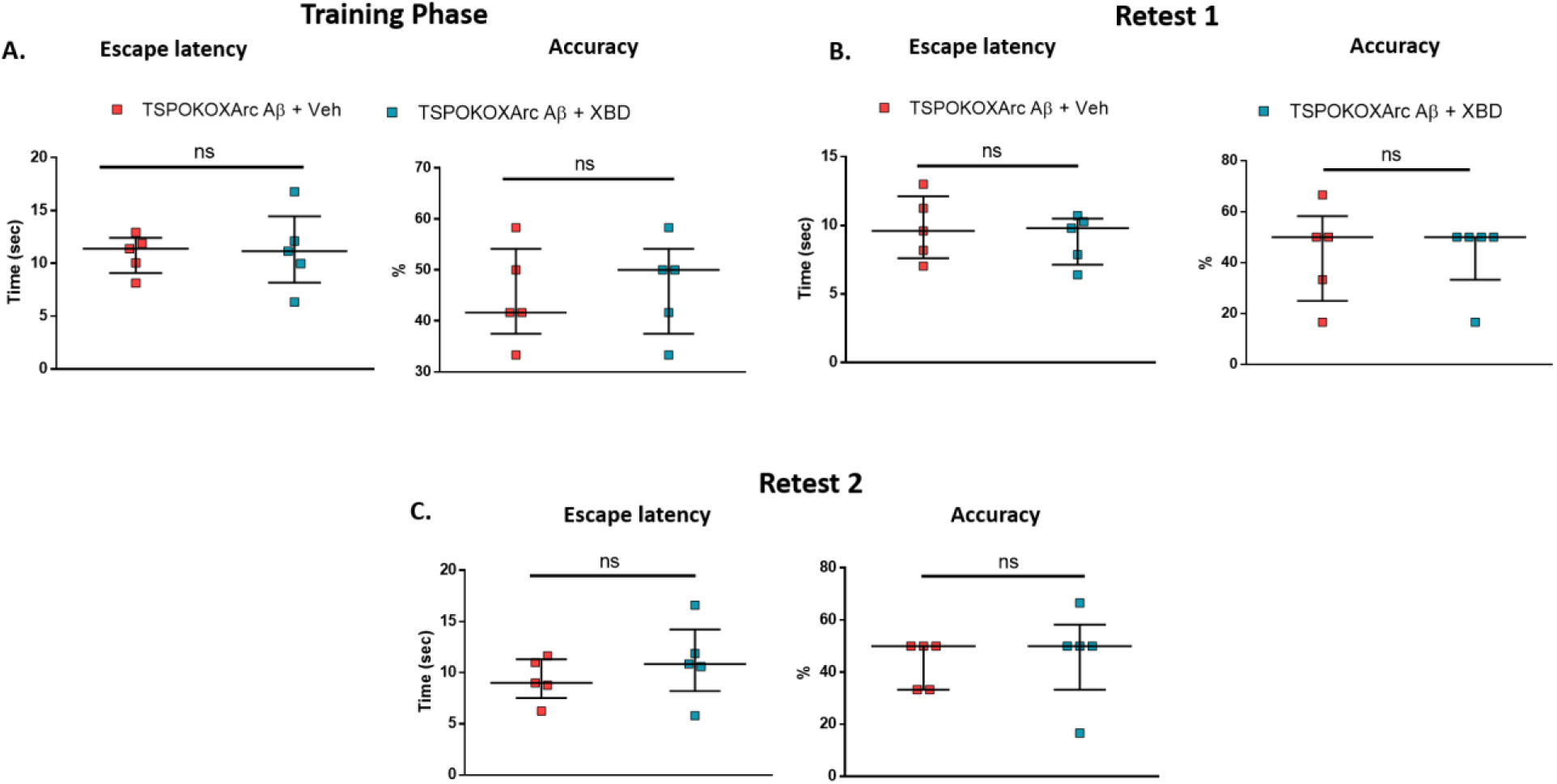
TSPO protein is responsible for XBD173-mediated spatial learning improvement. A. Escape latency (Mann–Whitney U test: hetTSPOKO X Arc Aβ + XBD: 11.16 (8.168- 14.46) s vs hetTSPOKO X Arc Aβ + Veh: 11.40 (9.097-12.43) s, *p=*0.944) and accuracy comparison (Mann–Whitney U test: hetTSPOKO X Arc Aβ + XBD: 50.00 (37.50-54.17) % vs hetTSPOKO X Arc Aβ + Veh (n=5): 41.67 (37.50-54.17) %, *p=*0.904) between the different treatment groups in the training phase. hetTSPOKO X Arc Aβ + Veh (n=5), and hetTSPOKO X Arc Aβ + XBD (n=5) groups. B. Escape latency (Mann–Whitney U test: hetTSPOKO X Arc Aβ + XBD: 9.800 (7.138-10.50) s vs hetTSPOKO X Arc Aβ + Veh: 9.600 (7.62-12.13) s, *p=*0.66) and accuracy comparison (Mann–Whitney U test: hetTSPOKO X Arc Aβ + XBD: 50.00 (33.33-50) % vs hetTSPOKO X Arc Aβ + Veh: 50.00 (25.00-58.33) %, *p>*0.99) between the different treatment groups in the Retest 1 phase. C. Escape latency (Mann–Whitney U test: hetTSPOKO X Arc Aβ + XBD: 10.86 (8.2-14.24) s vs hetTSPOKO X Arc Aβ + Veh: 9 (7.528-11.33) s, *p=*0.53) and accuracy comparison (Mann–Whitney U test: hetTSPOKO X Arc Aβ + XBD: 50.00 (33.33- 58.33) % vs hetTSPOKO X Arc Aβ + Veh: 50 (33.33-50) %, *p=*0.64) between the different treatment groups in the Retest 2 phase. Data are represented as median with their respective interquartile range. *\*p* < 0.05. ns: not significant.

**Supplementary Figure 9:**
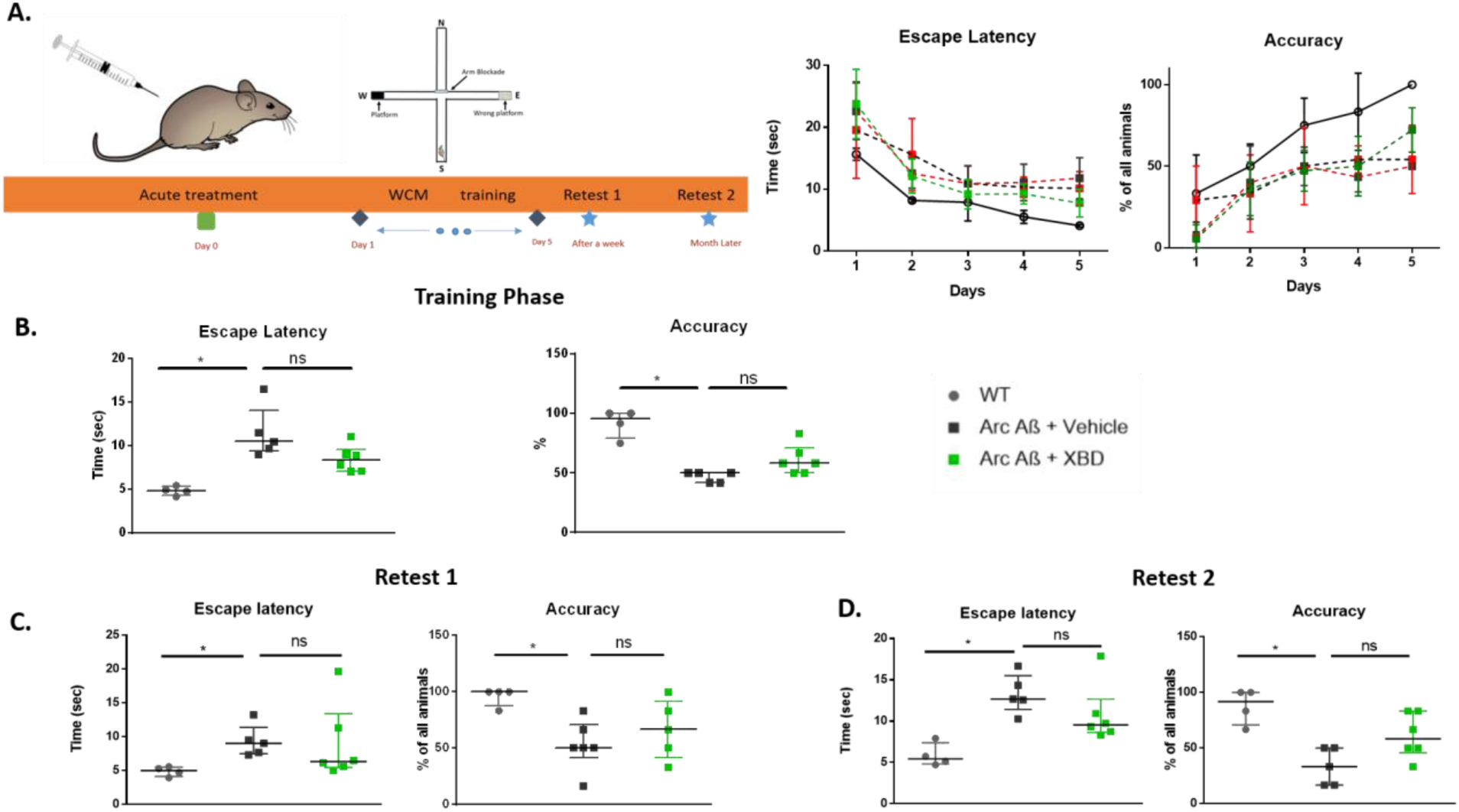
Acute administration of XBD173 doesn’t ameliorate cognitive deficits in AD mice. A. Schematic of acute treatment schedule, training, and retests in water cross maze. Adjacent to the schematic are the escape latency and accuracy curves during the course of the 5-day training phase in the water cross maze. B. Escape latency (Kruskal–Wallis test with a Dunn’s multiple comparisons *post hoc* test; Arc Aβ + vehicle: 10.47 (9.345-14.02) s vs Arc Aβ + XBD: 8.355 (7.095-9.548) s, *p=*0.27) and accuracy comparison (Kruskal–Wallis test with a Dunn’s multiple comparisons *post hoc* test; Arc Aβ + vehicle: 50 (41.67-50) % vs Arc Aβ + XBD: 58.33 (50-70.84) %, *p=*0.17) between the different treatment groups in the training phase. WT (n=4), Arc Aβ + vehicle (n=5), and Arc Aβ + XBD (n=6). Accuracy is expressed for each animal as the % of trials correctly performed, C. Escape latency (Kruskal–Wallis test with a Dunn’s multiple comparisons *post hoc* test; Arc Aβ + vehicle: 9.01 (7.473-11.36) s vs Arc Aβ + XBD: 6.302 (5.488- 13.39) s, *p=*0.69) and accuracy comparison (Kruskal–Wallis test with a Dunn’s multiple comparisons *post hoc* test; Arc Aβ + vehicle: 50 (41.67-70.83) % vs Arc Aβ + XBD: 66.66 (41.67-91.67) %, *p=*0.82) between the different treatment groups in the Retest 1 phase. D. Escape latency (Kruskal–Wallis test with a Dunn’s multiple comparisons *post hoc* test; Arc Aβ + vehicle: 12.67 (11.42-15.51) s vs Arc Aβ + XBD: 9.542 (8.643-12.67) s, *p=*0.44) and accuracy comparison (Kruskal–Wallis test with a Dunn’s multiple comparisons *post hoc* test; Arc Aβ + vehicle: 33.33 (16.66-50) % vs Arc Aβ + XBD: 58.33 (45.83-83.33) %, *p=*0.19) between the different treatment groups in the Retest 2 phase. Data are represented as median with their respective interquartile range. *\*p* < 0.05. ns: not significant.

**Supplementary Figure 10:**
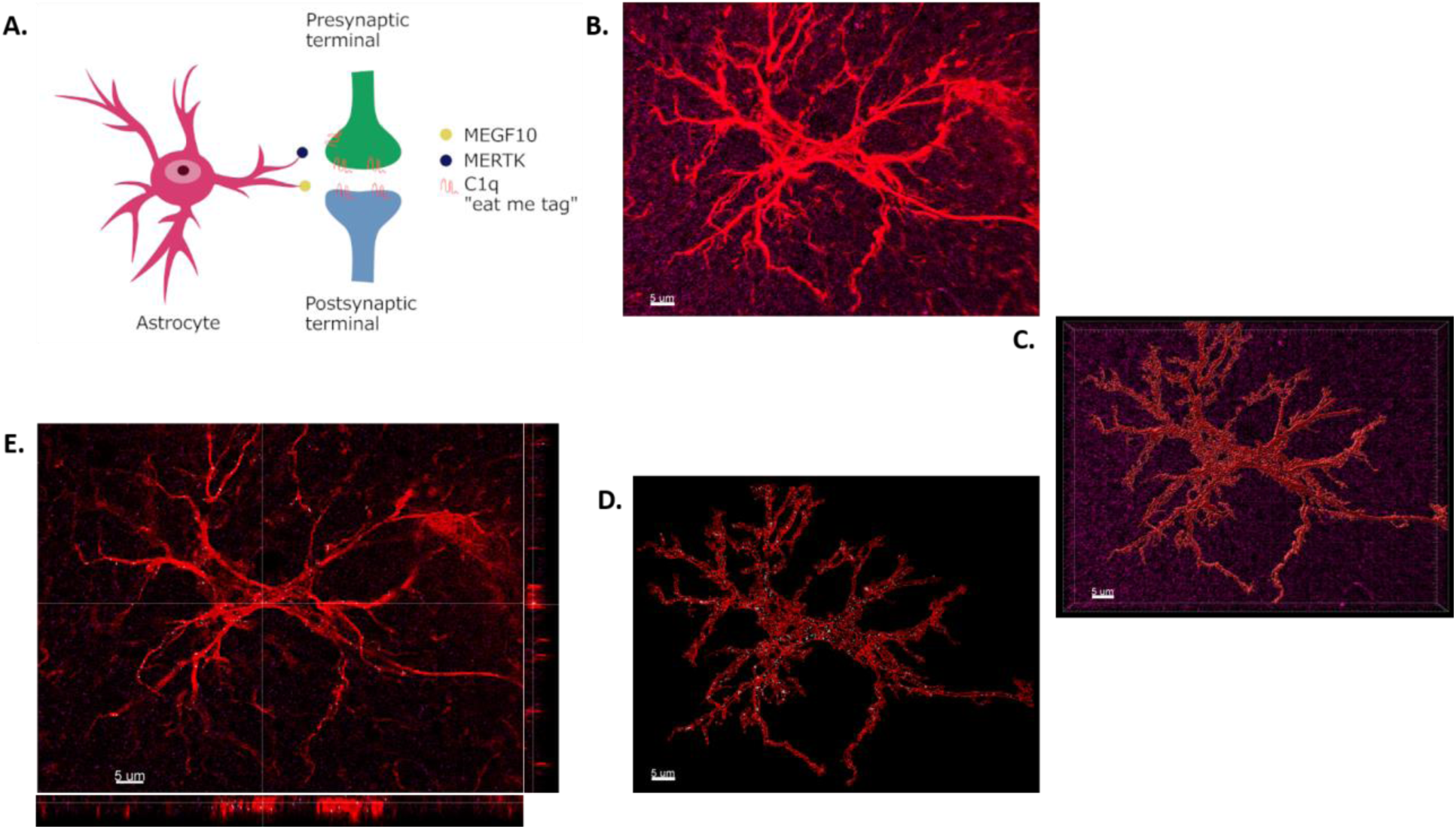
A. Schematic showing astrocytic receptors (MEGF10 and MERTK) in direct elimination of synapses via the C1q “eat-me tag”. B. Individual high-resolution astrocytes (shown in red) and C1q (shown in magenta). C. Volumetric reconstruction of astrocytes (red) using IMARIS 9.7 with C1q (magenta). D. Rendered outline of astrocyte (red) with colocalization between astrocytes and C1q marked in white. E. Orthogonal plane representation of astrocyte (red) along with colocalization (White). *Scale bar: 5 µm*.

## References

1. Rajmohan R, Reddy PH. Amyloid-beta and phosphorylated tau accumulations cause abnormalities at synapses of Alzheimer’s disease neurons. J Alzheimer’s Dis. 2017;57(4):975–99.

2. Tong L, Balazs R, Soiampornkul R, Thangnipon W, Cotman CW. Interleukin-1β impairs brain derived neurotrophic factor-induced signal transduction. Neurobiol Aging. 2008;29(9):1380–93.

3. Terrill-Usery SE, Mohan MJ, Nichols MR. Amyloid-β (1-42) protofibrils stimulate a quantum of secreted IL-1β despite significant intracellular IL-1β accumulation in microglia. Biochim Biophys Acta (BBA)-Molecular Basis Dis. 2014;1842(11):2276–85.

4. Zahs KR, Ashe KH. β-Amyloid oligomers in aging and Alzheimer’s disease. Front Aging Neurosci. 2013;5:28.

5. Walsh DM, Selkoe DJ. Deciphering the molecular basis of memory failure in Alzheimer’s disease. Neuron. 2004;44(1):181–93.

6. Villalobos Acosta DMÁ, Chimal Vega B, Correa Basurto J, Fragoso Morales LG, Rosales Hernández MC. Recent advances by in silico and in vitro studies of amyloid-β 1-42 fibril depicted a S-shape conformation. Int J Mol Sci. 2018;19(8):2415.

7. Zhang R, Hu X, Khant H, Ludtke SJ, Chiu W, Schmid MF, et al. Interprotofilament interactions between Alzheimer’s Aβ1–42 peptides in amyloid fibrils revealed by cryoEM. Proc Natl Acad Sci. 2009;106(12):4653–8.

8. Lei M, Xu H, Li Z, Wang Z, O’Malley TT, Zhang D, et al. Soluble Aβ oligomers impair hippocampal LTP by disrupting glutamatergic/GABAergic balance. Neurobiol Dis. 2016;85:111– 21.

9. Liu J, Chang L, Song Y, Li H, Wu Y. The role of NMDA receptors in Alzheimer’s disease. Front Neurosci. 2019;13:43.

10. Sáiz-Vázquez O, Gracia-García P, Ubillos-Landa S, Puente-Martínez A, Casado-Yusta S, Olaya B, et al. Depression as a risk factor for Alzheimer’s disease: a systematic review of longitudinal meta- analyses. J Clin Med. 2021;10(9):1809.

11. Cantón-Habas V, Rich-Ruiz M, Romero-Saldaña M, Carrera-González M del P. Depression as a risk factor for dementia and Alzheimer’s disease. Biomedicines. 2020;8(11):457.

12. Barnes DE, Yaffe K, Byers AL, McCormick M, Schaefer C, Whitmer RA. Midlife vs late-life depressive symptoms and risk of dementia: differential effects for Alzheimer disease and vascular dementia. Arch Gen Psychiatry. 2012;69(5):493–8.

13. Rupprecht R, Papadopoulos V, Rammes G, Baghai TC, Fan J, Akula N, et al. Translocator protein (18 kDa)(TSPO) as a therapeutic target for neurological and psychiatric disorders. Nat Rev Drug Discov. 2010;9(12):971–88.

14. Corsi F, Baglini E, Barresi E, Salerno S, Cerri C, Martini C, et al. Targeting TSPO Reduces Inflammation and Apoptosis in an In Vitro Photoreceptor-Like Model of Retinal Degeneration. ACS Chem Neurosci. 2022;13(22):3188–97.

15. Smith SS, Shen H, Gong QH, Zhou X. Neurosteroid regulation of GABAA receptors: focus on the α4 and δ subunits. Pharmacol Ther. 2007;116(1):58–76.

16. Tournier BB, Tsartsalis S, Ceyzeriat K, Fraser BH, Grégoire M-C, Kövari E, et al. Astrocytic TSPO upregulation appears before microglial TSPO in Alzheimer’s disease. J Alzheimer’s Dis. 2020;77(3):1043–56.

17. Rupprecht R, Rammes G, Eser D, Baghai TC, Schüle C, Nothdurfter C, et al. Translocator protein (18 kD) as target for anxiolytics without benzodiazepine-like side effects. Science (80-). 2009;325(5939):490–3.

18. Mihalek RM, Banerjee PK, Korpi ER, Quinlan JJ, Firestone LL, Mi Z-P, et al. Attenuated sensitivity to neuroactive steroids in γ-aminobutyrate type A receptor delta subunit knockout mice. Proc Natl Acad Sci. 1999;96(22):12905–10.

19. Rammes G, Seeser F, Mattusch K, Zhu K, Haas L, Kummer M, et al. The NMDA receptor antagonist Radiprodil reverses the synaptotoxic effects of different amyloid-beta (Aβ) species on long-term potentiation (LTP). Neuropharmacology. 2018;140:184–92.

20. Kleinknecht KR, Bedenk BT, Kaltwasser SF, Grünecker B, Yen Y-C, Czisch M, et al. Hippocampus-dependent place learning enables spatial flexibility in C57BL6/N mice. Front Behav Neurosci. 2012;6:87.

21. Pradhan AK, Shi Q, Tartler KJ, Rammes G. Quantification of astrocytic synaptic pruning in mouse hippocampal slices in response to ex vivo Aβ treatment via colocalization analysis with C1q. STAR Protoc. 2022;3(4):101687.

22. Schindelin J, Arganda-Carreras I, Frise E, Kaynig V, Longair M, Pietzsch T, et al. Fiji: an open- source platform for biological-image analysis. Nat Methods. 2012;9(7):676–82.

23. Zhu X, Fréchou M, Liere P, Zhang S, Pianos A, Fernandez N, et al. A role of endogenous progesterone in stroke cerebroprotection revealed by the neural-specific deletion of its intracellular receptors. J Neurosci. 2017;37(45):10998–1020.

24. Villers A, Giese KP, Ris L. Long-term potentiation can be induced in the CA1 region of hippocampus in the absence of αCaMKII T286-autophosphorylation. Learn Mem. 2014;21(11):616–26.

25. Malenka RC, Bear MF. LTP and LTD: an embarrassment of riches. Neuron. 2004;44(1):5–21.

26. Knobloch M, Konietzko U, Krebs DC, Nitsch RM. Intracellular Aβ and cognitive deficits precede β-amyloid deposition in transgenic arcAβ mice. Neurobiol Aging. 2007;28(9):1297–306.

27. Guo S, Wang H, Yin Y. Microglia polarization from m1 to m2 in neurodegenerative diseases. Front Aging Neurosci. 2022;14.

28. Wang Q, Yao H, Liu W, Ya B, Cheng H, Xing Z, et al. Microglia polarization in Alzheimer’s disease: mechanisms and a potential therapeutic target. Front Aging Neurosci. 2021;13.

29. Dejanovic B, Wu T, Tsai M-C, Graykowski D, Gandham VD, Rose CM, et al. Complement C1q- dependent excitatory and inhibitory synapse elimination by astrocytes and microglia in Alzheimer’s disease mouse models. Nat Aging. 2022;2(9):837–50.

30. Papadopoulos V, Aghazadeh Y, Fan J, Campioli E, Zirkin B, Midzak A. Translocator protein- mediated pharmacology of cholesterol transport and steroidogenesis. Mol Cell Endocrinol. 2015;408:90–8.

31. Rupprecht R, Wetzel CH, Dorostkar M, Herms J, Albert NL, Schwarzbach J, et al. Translocator protein (18kDa) TSPO: a new diagnostic or therapeutic target for stress-related disorders? Mol Psychiatry. 2022;1–9.

32. Lejri I, Grimm A, Hallé F, Abarghaz M, Klein C, Maitre M, et al. TSPO ligands boost mitochondrial function and pregnenolone synthesis. J Alzheimer’s Dis. 2019;72(4):1045–58.

33. Da Pozzo E, Giacomelli C, Costa B, Cavallini C, Taliani S, Barresi E, et al. TSPO PIGA ligands promote neurosteroidogenesis and human astrocyte well-being. Int J Mol Sci. 2016;17(7):1028.

34. Morlock E V, Czajkowski C. Different residues in the GABAA receptor benzodiazepine binding pocket mediate benzodiazepine efficacy and binding. Mol Pharmacol. 2011;80(1):14–22.

35. Dimitrova-Shumkovska J, Krstanoski L, Veenman L. Diagnostic and therapeutic potential of TSPO studies regarding neurodegenerative diseases, psychiatric disorders, alcohol use disorders, traumatic brain injury, and stroke: An update. Cells. 2020;9(4):870.

36. Pchitskaya E, Bezprozvanny I. Dendritic spines shape analysis—Classification or clusterization? Perspective. Front Synaptic Neurosci. 2020;12:31.

37. Shi Y, Cui M, Ochs K, Brendel M, Strübing FL, Briel N, et al. Long-term diazepam treatment enhances microglial spine engulfment and impairs cognitive performance via the mitochondrial 18 kDa translocator protein (TSPO). Nat Neurosci. 2022;25(3):317–29.

38. Batarseh A, Papadopoulos V. Regulation of translocator protein 18 kDa (TSPO) expression in health and disease states. Mol Cell Endocrinol. 2010;327(1–2):1–12.

39. Nutma E, Ceyzériat K, Amor S, Tsartsalis S, Millet P, Owen DR, et al. Cellular sources of TSPO expression in healthy and diseased brain. Eur J Nucl Med Mol Imaging. 2021;1–18.

40. Parsons CG, Stöffler A, Danysz W. Memantine: a NMDA receptor antagonist that improves memory by restoration of homeostasis in the glutamatergic system-too little activation is bad, too much is even worse. Neuropharmacology. 2007;53(6):699–723.

41. Griffin CE, Kaye AM, Bueno FR, Kaye AD. Benzodiazepine pharmacology and central nervous system–mediated effects. Ochsner J. 2013;13(2):214–23.

42. Inagaki T, Miyaoka T, Tsuji S, Inami Y, Nishida A, Horiguchi J. Adverse reactions to zolpidem: case reports and a review of the literature. Prim Care Companion CNS Disord. 2010;12(6):26600.

43. Baldwin DS, Aitchison K, Bateson A, Curran HV, Davies S, Leonard B, et al. Benzodiazepines: risks and benefits. A reconsideration. J Psychopharmacol. 2013;27(11):967–71.

44. Uzun S, Kozumplik O, Jakovljević M, Sedić B. Side effects of treatment with benzodiazepines. Psychiatr Danub. 2010;22(1):90–3.

45. Puig-Bosch X, Bieletzki S, Zeilhofer HU, Rudolph U, Antkowiak B, Rammes G. Midazolam at low nanomolar concentrations affects long-term potentiation and synaptic transmission predominantly via the α1–γ-aminobutyric acid type A receptor subunit in mice. Anesthesiology. 2022;136(6):954–69.

46. Whittington RA, Virág L, Gratuze M, Lewkowitz-Shpuntoff H, Cheheltanan M, Petry F, et al. Administration of the benzodiazepine midazolam increases tau phosphorylation in the mouse brain. Neurobiol Aging. 2019;75:11–24.

47. Owen DR, Guo Q, Kalk NJ, Colasanti A, Kalogiannopoulou D, Dimber R, et al. Determination of [11C] PBR28 binding potential in vivo: a first human TSPO blocking study. J Cereb Blood Flow Metab. 2014;34(6):989–94.

48. Mages K, Grassmann F, Jägle H, Rupprecht R, Weber BHF, Hauck SM, et al. The agonistic TSPO ligand XBD173 attenuates the glial response thereby protecting inner retinal neurons in a murine model of retinal ischemia. J Neuroinflammation. 2019;16(1):1–16.

49. Lee SY, Chung W-S. The roles of astrocytic phagocytosis in maintaining homeostasis of brains. J Pharmacol Sci. 2021;145(3):223–7.

50. Verkhratsky A, Parpura V, Rodriguez-Arellano JJ, Zorec R. Astroglia in Alzheimer’s disease. Neuroglia Neurodegener Dis. 2019;273–324.

51. Wu T, Dejanovic B, Gandham VD, Gogineni A, Edmonds R, Schauer S, et al. Complement C3 is activated in human AD brain and is required for neurodegeneration in mouse models of amyloidosis and tauopathy. Cell Rep. 2019;28(8):2111–23.

52. Rupprecht C, Rupprecht R, Rammes G. C1q, a small molecule with high impact on brain development: putative role for aging processes and the occurrence of Alzheimer’s disease. Vol. 271, European Archives of Psychiatry and Clinical Neuroscience. Springer; 2021. p. 809–12.

53. Rainer Rupprecht, Arpit Kumar Pradhan, Marco Kufner, Lisa Marie Brunner, Caroline Nothdurfter, Simon Wein, Jens Schwarzbach, Xenia Puig, Christian Rupprecht GR. Neurosteroids, translocator protein 18 kDa (TSPO) in depression: implications for synaptic plasticity, cognition, and treatment options. Eur Arch Psychiatry Clin Neurosci. 2022;In press.

54. Karlstetter M, Nothdurfter C, Aslanidis A, Moeller K, Horn F, Scholz R, et al. Translocator protein (18 kDa)(TSPO) is expressed in reactive retinal microglia and modulates microglial inflammation and phagocytosis. J Neuroinflammation. 2014;11(1):1–13.

55. Hulshof LA, Van Nuijs D, Hol EM, Middeldorp J. The Role of Astrocytes in Synapse Loss in Alzheimer’s Disease: A Systematic Review. Front Cell Neurosci. 2022;16:899251.

56. Hong S, Beja-Glasser VF, Nfonoyim BM, Frouin A, Li S, Ramakrishnan S, et al. Complement and microglia mediate early synapse loss in Alzheimer mouse models. Science (80-). 2016;352(6286):712–6.

57. Henstridge CM, Tzioras M, Paolicelli RC. Glial contribution to excitatory and inhibitory synapse loss in neurodegeneration. Front Cell Neurosci. 2019;13:63.

58. Shah A, Kishore U, Shastri A. Complement System in Alzheimer’s Disease. Int J Mol Sci. 2021;22(24):13647.

59. Gomez-Arboledas A, Acharya MM, Tenner AJ. The role of complement in synaptic pruning and neurodegeneration. ImmunoTargets Ther. 2021;10:373.

60. Iram T, Ramirez-Ortiz Z, Byrne MH, Coleman UA, Kingery ND, Means TK, et al. Megf10 is a receptor for C1Q that mediates clearance of apoptotic cells by astrocytes. J Neurosci. 2016;36(19):5185–92.

61. Webster S, O’barr S, Rogers J. Enhanced aggregation and β structure of amyloid β peptide after coincubation with C1q. J Neurosci Res. 1994;39(4):448–56.

62. Sarvari M, Vago I, Weber CS, Nagy J, Gal P, Mak M, et al. Inhibition of C1q-β-amyloid binding protects hippocampal cells against complement mediated toxicity. J Neuroimmunol. 2003;137(1– 2):12–8.

63. Carpanini SM, Torvell M, Bevan RJ, Byrne RAJ, Daskoulidou N, Saito T, et al. Terminal complement pathway activation drives synaptic loss in Alzheimer’s disease models. Acta Neuropathol Commun. 2022;10(1):1–16.

64. Johnson L V, Leitner WP, Rivest AJ, Staples MK, Radeke MJ, Anderson DH. The Alzheimer’s Aβ- peptide is deposited at sites of complement activation in pathologic deposits associated with aging and age-related macular degeneration. Proc Natl Acad Sci. 2002;99(18):11830–5.

65. Tenner AJ. Complement-mediated events in Alzheimer’s disease: mechanisms and potential therapeutic targets. J Immunol. 2020;204(2):306–15.

66. Ravikumar B, Crawford D, Dellovade T, Savinainen A, Graham D, Liere P, et al. Differential efficacy of the TSPO ligands etifoxine and XBD-173 in two rodent models of Multiple Sclerosis. Neuropharmacology. 2016;108:229–37.

67. Wang M. Neurosteroids and GABA-A receptor function. Front Endocrinol (Lausanne). 2011;2:44.

68. Brickley SG, Mody I. Extrasynaptic GABAA receptors: their function in the CNS and implications for disease. Neuron. 2012;73(1):23–34.

69. 69. Kennedy A. mouse profile. 2020 Jul 1 [cited 2023 Feb 16]; Available from: https://zenodo.org/record/3925921

70. Simon W, Hapfelmeier G, Kochs E, Zieglgänsberger W, Rammes G. Isoflurane blocks synaptic plasticity in the mouse hippocampus. J Am Soc Anesthesiol. 2001;94(6):1058–65.

